# Stimulus and circuit contributions to the information geometry of neural manifolds

**DOI:** 10.64898/2026.06.21.733384

**Authors:** Sven Goedeke, Justus Kautz, Christian Leibold

## Abstract

Understanding how network connectivity shapes neural representations is central to systems neuroscience. While dimensionality reduction methods uncover low-dimensional manifold structure in population recordings, a rigorous framework connecting manifold geometry to network mechanisms and information encoding remains lacking. We develop a differential geometric approach for analyzing neural manifolds in rate-based recurrent networks receiving tuned feedforward inputs. We derive expressions for the pullback metric of neural manifolds, showing how input tuning curves together with feedforward and recurrent synaptic connectivity shape manifold geometry. Critically, we establish that the Fisher information matrix at steady states also has the structure of a pullback metric, directly linking intrinsic manifold geometry to stimulus discriminability and information encoding. For noise with slow temporal correlations propagated through the network, we show that recurrent effects on information geometry cancel: Fisher information depends only on the feedforward connectivity. Thus, feedforward connectivity critically determines representational geometry. We apply our approach to the representation of space by a module of hexagonal grid cells. We first demonstrate that the representation is approximately isometric for a random distribution of grid phases. Moreover, a linear feedforward transformation can map spatially random input tuning curves into a population of hexagonal grid cells, generating a toroidal neural manifold. Thus, feedforward connectivity alone can generate structured spatial representations without requiring carefully tuned recurrent connectivity or continuous attractor dynamics. Recurrent connectivity, however, is shown to improve stimulus encoding under fast noise, thereby implementing a selective noise reduction.

## I. INTRODUCTION

Information processing in biological and artificial neural networks is based on interactions of populations of neurons. Many successful approaches to decipher neural population codes take the perspective of a single readout layer, which led to the development of theories grounded in the statistical physics of linear separability [1–4]. The success of deep neural networks [5, 6], however, showed that complex nonlinear structures of stimulus representations can be revealed several synapses further downstream. Nonlinear structures in neural activity spaces may therefore represent meaningful information, but cannot be directly accessed with linear approaches.

Triggered by the development of modern recording techniques, such as high-density neuropixels probes [7] and population calcium imaging [8], which enabled simultaneous recording from hundreds of neurons in behaving animals, low-dimensional, typically nonlinear structures of neural activity are increasingly characterized as neural manifolds [9–12]. While the manifold approach has generated important insights, particularly the discovery that neural embedding spaces contain low-dimensional structures that capture behaviorally relevant parameters, sensory representations, or cognitive variables [9, 13, 14], so far, no consensus has been reached on how the neural population code on these nonlinear structures should be quantified (but see [15, 16]). In particular, neural manifolds are often only interpreted from a topological point of view [13, 14], whereas geometric aspects of manifolds are considered only in certain aspects [16, 17]. Furthermore, the connection between manifold topology/geometry and the underlying neural circuit mechanisms is still rather correlative [10], missing more fundamental theoretical consideration.

In this paper, we argue that the existing theory of information geometry [18, 19] is not only inherently designed to deal with nonlinear coding problems and well-suited to understand the neural manifold idea more thoroughly [15], it also provides a clear and rigorous interpretation of geometry, as well as a connection to circuit parameters and model assumptions. We focus our approach on attractor states in linear networks but point towards possible generalizations. In illustrative examples, we demonstrate that even in linear feedforward networks, distortions of geometry, as well as changes in topology, can be realized. We show how the effect of recurrent synaptic connections crucially depends on the underlying noise assumptions [20]. Finally, we show how our formalism can be employed to provide analytical results on the isometry of representations of space in medial entorhinal grid cell modules.

## II. THEORY

### A. Model Setup

We base our considerations on rate-based neural networks where the dynamics of neuron *i* is governed by

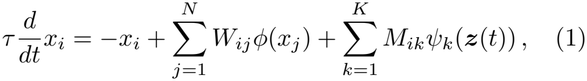

for *i* = 1, …, *N* . Here, *x*_*i*_ is the membrane potential (preactivation), *τ >* 0 is the membrane time constant, and *ϕ* : ℝ→ ℝis a generally nonlinear differentiable activation function. The matrix *W* ∈ ℝ^*N ×N*^ contains the recurrent connection weights in the network, *M* ∈ ℝ^*N ×K*^ is a matrix of feedforward input weights to the network, and for each input neuron *k*, the function *ψ*_*k*_(***z***) yields the tuning of this input neuron to some stimulus variable ***z***, which could either represent a sensory stimulus or, more generally, latent features of inputs from another neuron population. We write the network dynamics in vector-matrix form as

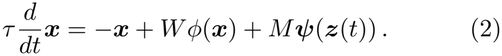

Within this paper, we will only consider steady state solutions (fixed points) ***x***^*\**^ ∈ ℝ^*N*^ of Eq. (2), thereby assuming that the stimulus ***z***(*t*) varies slowly compared to the network dynamics. For a constant stimulus value ***z***, a steady state solves the implicit equation

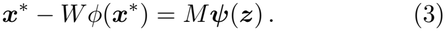

We will focus on the important case, where the steady state ***x***^*\**^(***z***) is the unique fixed-point attractor of Eq. (2). In particular, we assume that the linearized dynamics near the fixed point ***x***^*\**^ is stable, i.e. all eigenvalues of the Jacobian matrix 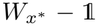 have negative real parts, where 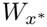 denotes the matrix with entries 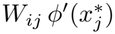.

To obtain the derivative (Jacobian matrix) of ***x***^*\**^(***z***) with respect to ***z*** (coordinates of the stimulus), we differentiate Eq. (3):

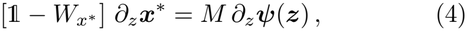

where *∂*_*z*_ indicates the row vector of partial derivatives with respect to the coordinates of ***z***, applied elementwise to ***x***^*\**^ and ***ψ***. With our assumption of a stable fixed point, the matrix 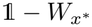 is invertible and we can solve Eq. (4) for the derivative *∂*_*z*_***x***^*\**^(***z***). To simplify notation, we will drop the ***x***^*\**^-dependence of 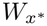 in the following, as all example applications will use the linear activation function *ϕ*(*x*) = *x*.

### B. Manifolds

As a *stimulus manifold* we consider a Riemannian manifold (*Ƶ, g* _*Ƶ*_), with metric *g* _*Ƶ*_, spanned by all variables that determine the input to a brain area, e.g., position, intensity, head direction, etc. Stimuli ***z*** ∈ *Ƶ* evoke neural activity according to tuning curves *ψ*_*k*_ : *Ƶ* → ℝ, ***z*** ↦ *ψ*_*k*_(***z***), which are usually nonlinear differentiable functions. An example we use in the first application are Gaussian tuning functions

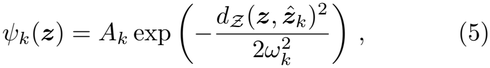

with 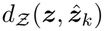 denoting the geodesic distance between ***z*** and a reference point 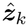 in the stimulus manifold (stimulus preference). For a population of *K* input neurons with tuning curves *ψ*_*k*_(***z***), the image of the stimulus manifold, *J* = ***ψ*** (*Ƶ*) ⊂ ℝ^*K*^, is then called the *input manifold*, with the standard Euclidean inner product in ℝ^*K*^ inducing its metric (Sec. II C).

By varying the stimulus ***z*** quasi-statically, the steady state ***x***^*\**^(***z***) of the network dynamics, Eq. (2), traces out a set of points in the activity space ℝ^*N*^ . We consider this set ***x***^*\**^(*Ƶ*) as the *neural manifold ℳ* embedded in the ambient space ℝ^*N*^, in which we again use the standard Euclidean inner product.

### C. Representational Geometry

The geometry of a manifold is governed by its metric tensor [21–23], which allows one to measure distances and volumes on the manifold and angles in its tangent space. The metric tensor thus sufficiently describes all geometric properties of processes that are constrained to the manifold, i.e., those that have no information about the embedding space orthogonal to the tangent space. The metric tensor therefore characterizes the intrinsic geometry, in contrast to the extrinsic geometry, which describes how a manifold is shaped in the embedding space (for relevance of extrinsic geometry, see Discussion).

The input and neural manifold inherit the metric from the respective embedding space, i.e., the standard Euclidean inner product. These metrics can therefore be “pulled back” to the stimulus space as follows. For the input manifold *J*, we obtain

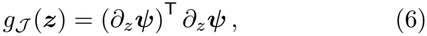

For the neural manifold *ℳ*, we use Eq. (4) to find

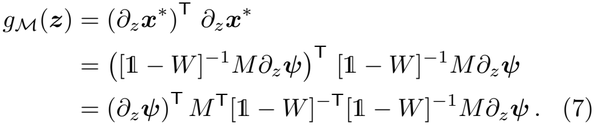

The intrinsic geometry of the neural manifold thus combines effects of recurrent connections *W*, feedforward connections *M*, and the deformation introduced by the nonlinear tuning functions.

Having pulled back the metrics of the input and the neural manifold to the stimulus space, we can visualize them as functions of ***z*** and compare them with the metric *g* _*Ƶ*_ . The pullback metric quantifies lengths and angles on the manifold in a certain region of the stimulus space. Intuitively, an increase or decrease in length leads to an increase or decrease in resolution and stimulus discriminability. The synaptic weight matrices may be able to warp regions around decision boundaries in downstream areas for better discriminability. Relevant regions in stimulus space could be selectively expanded while less relevant regions could be compressed [24].

The intuition derived from stimulus encoding/representational geometry is in line with our basic understanding of decoding limits induced by local discriminability, i.e., Fisher information and the Cramér-Rao bound [15]. However, these statistical concepts, at first glance, do not directly apply to a deterministic steady state ***x***^*\**^(***z***), Eq. (3). We therefore assume that the observed activity ***x*** varies around its expectation value ***x***^*\**^(***z***),

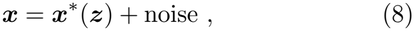

according to a noise distribution *p*(***x*** | ***z***). The correlation structure of the noise adds further degrees of complexity to stimulus discriminability, which is classically treated via information geometry [18], the relation of which to neural manifolds we will lay out in the following.

### D Information Geometry and Neural Manifolds

The Fisher information matrix (sensitivity matrix) is derived from the noise distribution *p*(***x***|***z***) as

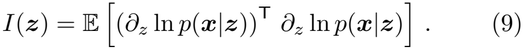

It provides an asymptotic lower bound for the mean squared error *ℰ* of an unbiased estimator of ***z*** via the Cramér-Rao bound [25]

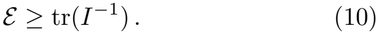

The Fisher information matrix is symmetric and positive semi-definite. Therefore, if it has full rank, it qualifies as a metric tensor of the *statistical manifold* formed by the family of probability distributions *p*(***x***| ***z***) [18], allowing to measure distances between stimuli ***z***∈ *Ƶ* based on their statistical separability. To relate the statistical manifold and its metric *I*(***z***) to the neural manifold *ℳ* and enable mechanistic interpretations in terms of neural networks and synaptic mechanisms, we assume that the noise distribution depends on the stimulus space *Ƶ* only via the expectation value (mean activity) ***x***^*\**^(***z***), i.e.,

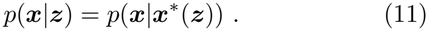

This excludes the case that two different stimuli ***z*** yield the same ***x***^*\**^(***z***) but different noise distributions, in which only the statistical manifold but not the neural manifold bears significance for the neural code. This property is fundamental to the neural manifold idea that defines manifolds in the space of firing rates.

Under this assumption, the Fisher information from Eq. (9) transforms as

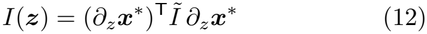

with the symmetric and positive (semi-)definite matrix *Ĩ*(***x***^*\**^) ∈ ℝ^*N ×N*^ denoting the Fisher information of the noise distribution *p*(***x*** |***x***^*\**^) on the neural manifold ***x***^*\**^(***z***). We note that this expression has the same form as the linear Fisher information [20] where *Ĩ* is replaced by the inverse covariance matrix of the noise distribution.

Since *Ĩ* is positive semidefinite, a square root *Ĩ*^1*/*2^ with *Ĩ* = (*Ĩ*^1*/*2^)^⊤^*Ĩ*^1*/*2^ exists. Thus, a manifold Y = *y*(*Ƶ*) (the statistical manifold) that has its tangent space spanned by the Jacobian matrix *∂*_*z*_ ***y*** = *Ĩ* ^1*/*2^*∂*_*z*_***x***^*\**^ has the Fisher information (12) as its metric tensor induced from the Euclidean inner product:

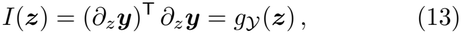

which has the same algebraic form as *g*_*ℳ*_ (***z***) above, except for ***x***^*\**^ replaced by ***y***. As an illustration, we next consider a few straight-forward but important examples.

#### Example 1: Stimulus-Independent Isotropic Gaussian Noise

In the special case of an isotropic Gaussian noise with covariance matrix *Σ*^2^1 that is independent of the stimulus, we find *Ĩ* = *Σ*^*Δ*2^ 1 and

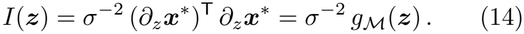

Thus, the Fisher information *I*(***z***) is proportional to the pullback metric *g* _*ℳ*_ (***z***) of the neural manifold. In this case (which will be mostly discussed in the next section), neural manifolds match the intuition of providing a mechanistic account for the information geometry of neuronal rate networks.

#### Example 2: Non-Isotropic Gaussian Noise

In the case of Gaussian noise with general (invertible) covariance matrix Σ, we find the Fisher information

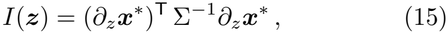

and thus *Ĩ* = Σ^*Δ*1^. In the case of stimulus-dependent covariance Σ = Σ(***z***), the Fisher information contains an additional “trace-term”, so that

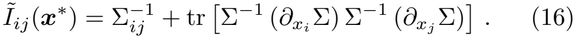

In the applications of Sec. III, we will generally use Eq. (15) for simplicity but keep in mind that Σ^*Δ*1^ can be replaced by more general *Ĩ*(***x***). A simple example provides a diagonal covariance with entries Σ_*ii*_ that depend only on the corresponding variable *x*_*i*_, i.e.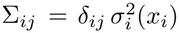. Then, the matrix *Ĩ* is diagonal as well and

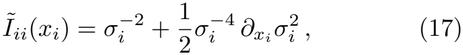

which generalizes the case of stimulus-independent isotropic Gaussian noise (*Example 1*).

If the noise covariance Σ is independent of ***x***^*\**^(***z***), the transformation between the neural and the statistical manifold is simply linear,

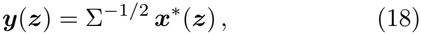

with the above-mentioned square root of Σ^*Δ*1^ = *Ĩ*. Thus activity dimensions with high noise will be compressed and directions with low noise will be expanded. In the case of stimulus-dependent Σ(***z***), the Jacobian of the statistical manifold with ***y***(***z***) satisfies

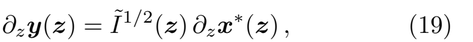

corresponding to local whitening. This local whitening can be realized by a nonlinear transformation ***y*** = ***F*** (***x***^*\**^) if *Ĩ*^1*/*2^ satisfies the integrability conditions from upcoming Eq. (23).

##### Nonlinear Activity Transformations

Assuming there exists a differentiable function ***F*** (***x***^*\**^) such that ***y***(***z***) = ***F*** (***x***^*\**^(***z***)), the tangent vectors transform as

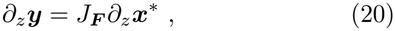

with *J*_***F***_ denoting the Jacobian of ***F*** . In order to satisfy the interpretation of the Fisher information as the metric of the statistic manifold according to Eq. (13), the Jacobian must obey

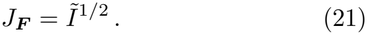

Such a function ***F*** exists, if the integrability conditions

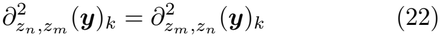

holds, i.e., if

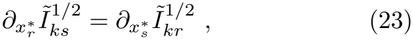

which is, e.g., always the case for constant *Ĩ*.

For diagonal 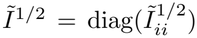 the integrability conditions (23) imply 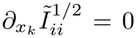 for *k* ≠= *i*, which means that 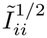 does only depend on *x*_*i*_. In this case, we can obtain the transformation ***F*** (***x***) by integrating Eq. (21) element-wise for each *i*:

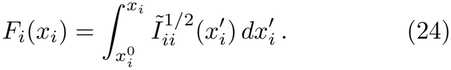

This approach will be illustrated with Poisson distributed activity in the next paragraph on *Poisson noise*.

Importantly, nonlinear transformations alter both the intrinsic geometry (distance matches statistical discriminability) and the shape (extrinsic geometry) of the manifold.

#### Example 3: Poisson Noise

For Poisson distributed spike counts ***x*** with mean ***x***^*\**^(***z***) during a time bin of width Δ*t*, the Fisher information equals

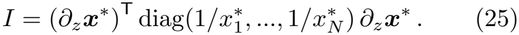

In this case, a nonlinear transformation ***F*** satisfying 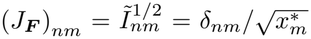 can be constructed according to Eq. (24) as

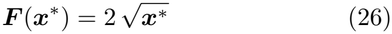

with the square root applied element-wise (known as the square-root transformation [26]). Owing to the ubiquitous assumption of Poisson firing in systems neuro-science, square-root transformation of neural activity has become a standard preprocessing procedure in neural manifold analyses [13, 14].

### E. Noise propagation

So far, the noise distribution *p*(***x*** |***x***^*\**^(***z***)) has been considered independent of the network dynamics. Fluctuations, however, may also be subject to the transformations imposed by the synaptic connectivity. We therefore assume that the deviations

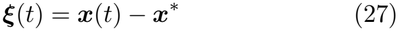

from the fixed point are driven by two sources of noise. Network noise ***η***(*t*) that is directly introduced in the neural activity space, and stimulus noise ***ζ***(*t*) that adds to the given stimulus ***z***. Both noise sources are assumed to be stationary Gaussian with covariance matrices Σ_*η*_(*t−t*^*′*^) = *E* ***η***(*t*)***η***(*t*^*′*^)^⊤^ and Σ_*ζ*_(*t−t*^*′*^), respectively. Assuming small noise amplitudes, we arrive at a differential equation for the activity fluctuations ***ξ***(*t*),

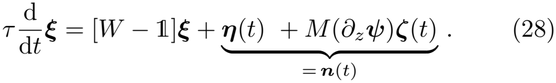

As a condition for the equal-time covariance matrix Σ_*ξ*_ = *E*[***ξ***(*t*) ***ξ***(*t*)^⊤^] of the activity to be stationary, 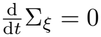, we obtain the continuous Lyapunov equation

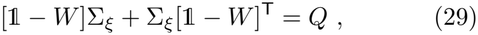

with effective noise covariance (App. A)

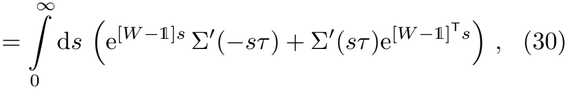

where Σ^*′*^ = Σ_*η*_ + *B*Σ_*ζ*_*B*^⊤^ and *B* = *M∂*_*z*_***ψ***. The general solution [27] of the continuous Lyapunov equation (29) is obtained as

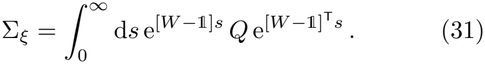

As a result, the Fisher information for the case that the noise is propagated by the network dynamics reads

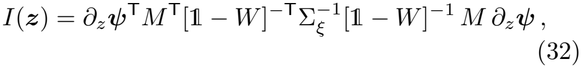

with the covariance matrix Σ_*ξ*_ from Eq. (31).

## III. APPLICATIONS

### A. Two neurons in low-rank network with Gaussian tuning

As a first illustrative example, we consider a 1-d stimulus ***z***∈ [Δ 1, 1] ⊂ℝ^1^ that is sampled by *K* = 2 Gaussian tuning curves

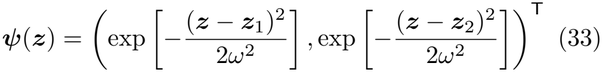

with centers located at ***z***_1_ = (Δ 0.5), ***z***_2_ = (+0.5) and tuning width *ω* = 0.75 (Fig. 1(a)).

**Figure 1.**
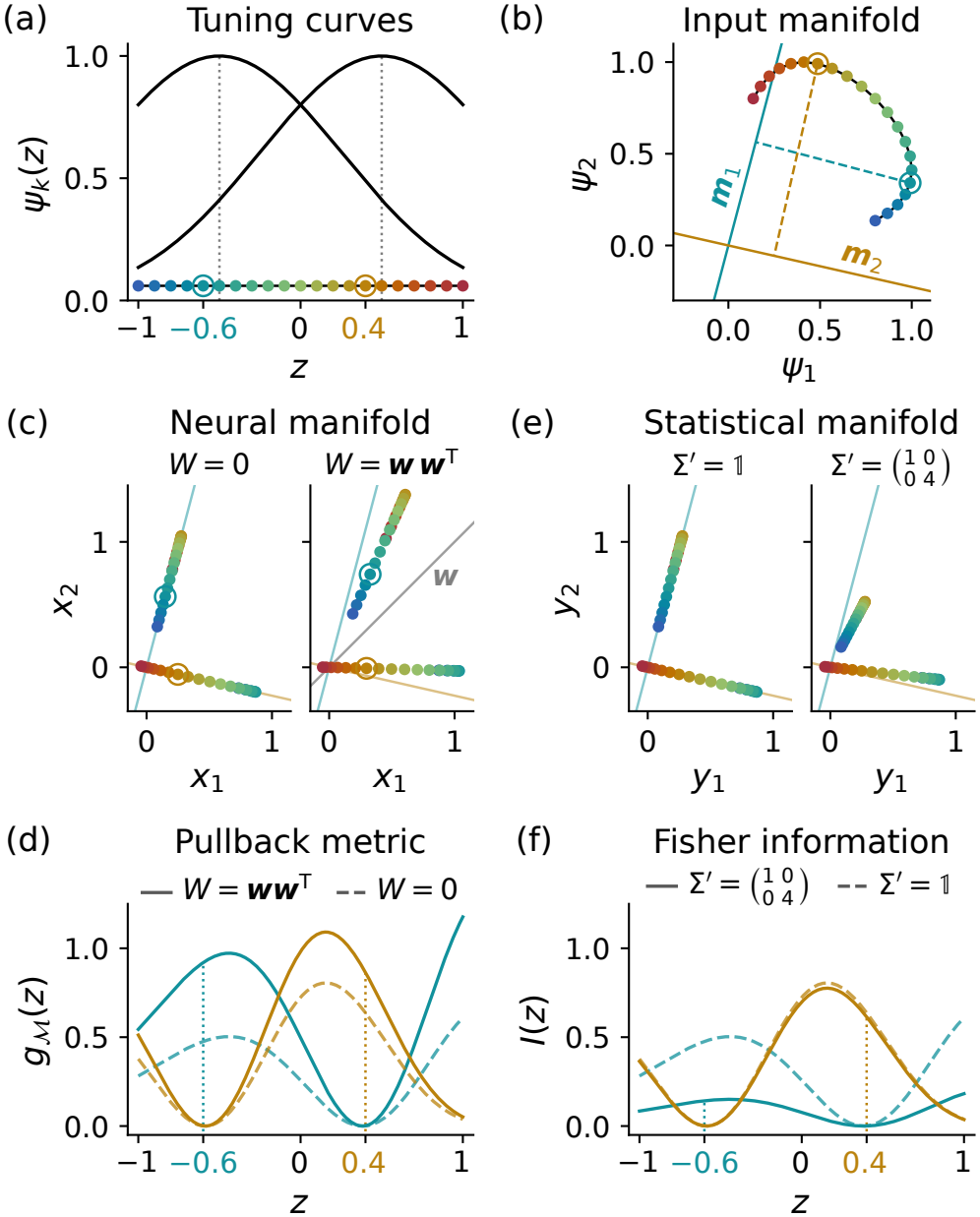
Manifolds in a two-neuron network. (a) Stimuli ***z*** are transformed into input activity by two tuning curves *ψ*_1_ and *ψ*_2_ (black). Equally spaced colored dots (blue to red) mark exemplary stimuli. Stimuli with circles will be focused on in the following panels. (b) The two tuning curves map the stimulus space into a 2-dimensional curve, the input manifold (black). Colored dots on the input manifold correspond to the stimuli from (a). Dots on the input manifold are no longer equidistant. Perpendicular projections (dashed) on 1-dimensional subspaces (solid lines) can be induced by the vector ***m*** that spans the feedforward subspace. For each stimulus ***z, m*** can be chosen parallel to the tangent space of the input manifold at ***z***. Two examples for ***z*** are shown (teal: ***m***_1_, orange: ***m***_2_) that correspond to the circles from (a). (c) The neural manifolds are the fixed points of the dynamics (teal for ***z*** = (−0.6) and orange for ***z*** = (0.4)). Left: ***w*** = 0. Due to the choice of ***m***, the distance (metric) between points on the neural manifold is particularly large for the corresponding stimulus. Adding low-rank recurrence parallel to ***w*** (right) can additionally expand the manifold along ***w***. (d) The metric reflects the distances between neighboring stimuli on the neural manifold. Colors correspond to the two choices of ***m*** from (b), dashed lines are derived from ***w*** = 0, solid lines for ***w*** = (1, 1)^T^. (e) For slow isotropic noise (left) the statistical manifold matches the neural manifold for ***w*** = 0. For higher noise in the *x*_2_-direction, the statistical manifold is squeezed along this noisy dimension. (f) Same as (d) for Fisher information. For slow noise, recurrent connectivity does not affect Fisher information (dashed lines are the same as in (d)). Increased noise along *x*_2_ direction selectively reduces Fisher information if ***m*** strongly aligns with *x*_2_ axis (teal).

As feedforward and recurrent weights we choose rank-1 projection matrices

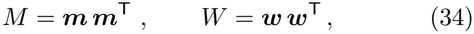

with ***w***^⊤^***w*** = | ***w***| ^2^ *<* 1 in order to preserve stability of the fixed point manifold

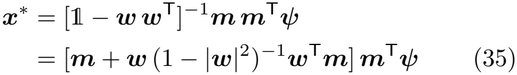

under the dynamics. The feedforward operation can be illustrated in the input space as a projection of the input manifold ***ψ***(***z***) onto the subspace spanned by ***m*** (Fig. 1(b)). Depending on the direction of ***m***, different parts of the input manifold are better resolved on the neural manifold (Fig. 1(c)). The recurrent weights only further expand the neural manifold along the direction ***w*** spanned by the recurrent matrix (Fig. 1(c), right). The pullback metric (Fig. 1(d)) reflects the warped geometry of the stimulus space, with large *g*_*ℳ*_ for stimuli ***z*** where neighbors have large distances on the manifold.

The choice of the “communication subspace” [28] of input and neural manifold spanned by ***m*** thus determines which stimuli ***z*** are best resolved. Recurrent weights can only scale the metric by a factor larger than one.

The transition from neural to statistical manifold requires computing the Fisher information (12), which for propagated Gaussian noise is given by Eq. (32). If we assume that the fluctuation covariance at the fixed-point is generated by the stable dynamics, Σ_*ξ*_ in Eq. (32) has to be computed from Eq. (31). For now, we only consider slow Gaussian input noise where Σ^*′*^(*t*) ≈ Σ^*′*^(0) = Σ^*′*^ describes its covariance over long time scales.

As shown in App. B for general diagonalizable *W*, the Fisher information from Eq. (32) for slow noise no longer depends on the recurrent weights and reads

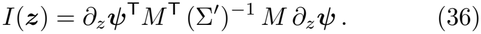

Intuitively, this is because *W* amplifies signal and noise equally. Thus, in our example with a 1-d stimulus ***z***, we get

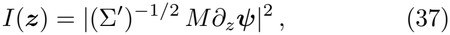

where | ***x***| ^2^ = ***x***^⊤^***x*** denotes the squared inner product of a vector ***x***.

In Fig. 1(e), we illustrate the statistical manifold for slow Gaussian noise, starting with the isotropic case with variance *Σ*^2^ = 1, so that Σ^*′*^ = 1 and *I*(*z*) = *g*_*ℳ*_(*z*) for *W* = 0. In this case, the statistical manifold is identical to the neural manifold without recurrent weights (Fig. 1(e), left). For non-isotropic noise, however, e.g., 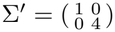, the statistical manifold is scaled with Σ^*′Δ*1*/*2^, which effectively compresses the high-noise direction, here the direction of *y*_2_ (Fig. 1(e), right, & (f)).

Therefore, if the slow noise is internal, in the sense that it is propagated through the network dynamics, recurrent weights cannot be chosen such that they compensate for the noise. In this case, recurrent connectivity, in contrast to feedforward connectivity, is unsuitable for improving stimulus representation.

As a next step, we evaluate these insights in a more complex situation with multiple input cells tuned to a circular stimulus.

### B. Representational geometry on a ring

To further illustrate transformations from stimulus geometry to input geometry and eventually geometry on the neural manifold, we consider a circular stimulus *z* ∈ [Δ *π, π*] (represented by the unit circle in Fig. 2(a)) and tuning curves *ψ*_*k*_(*z*), *k* = 1, …, *K*, of the input population as 2*π*-periodic functions, most parsimoniously,

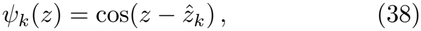

**Figure 2.**
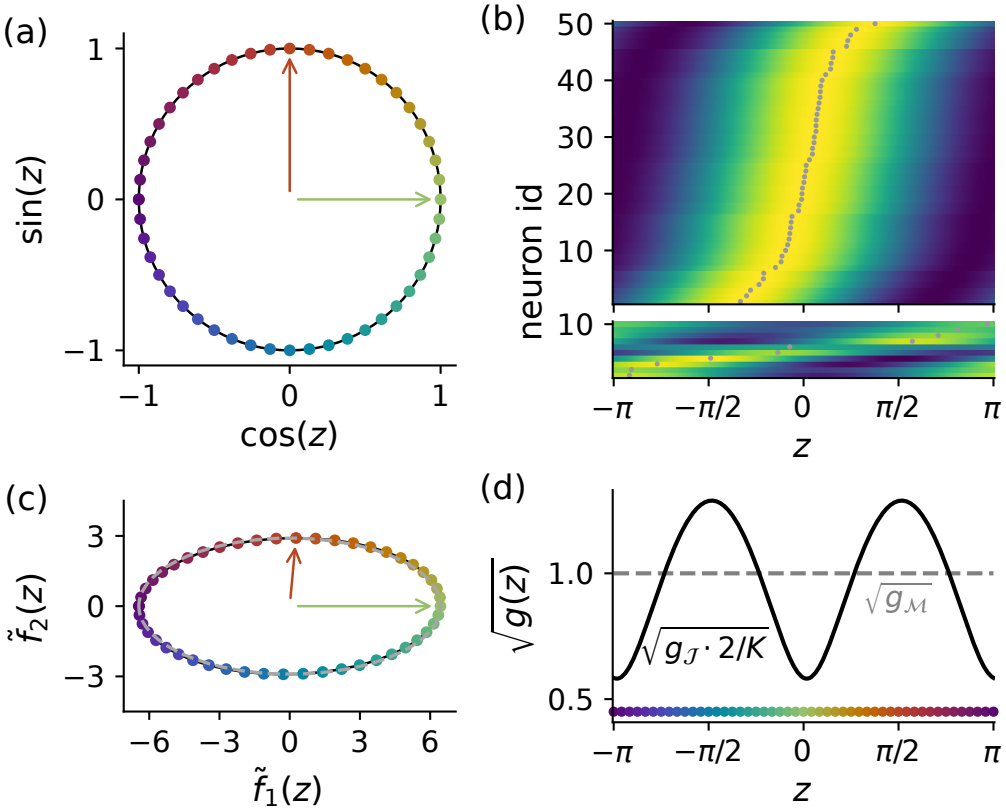
Ring geometry. (a) Latent representation of circular stimulus *z* ∈ [−*π, π*] as (cos(*z*), sin(*z*))^T^ ∈ ℝ. Colored dots mark equidistant values of *z*; the color code is indicated in panel (d). Arrows point to representations of *z* = 0 (green) and *z* = *π/*2 (red). (b) Tuning curves *ψ*_*k*_(*z*) of *K* = 50 input units (top) and activity *x*_*i*_(*z*) of *N* = 10 network neurons obtained via the feedforward transformation (bottom), each sorted by preferred stimulus (small gray dots). Color maps: top panel from − 1 (dark blue) to 1 (yellow); bottom panel symmetric around zero (dark blue to yellow). (c) Input manifold embedded in the two-dimensional subspace ofℝ spanned by the QR orthonormal basis of the input latent vectors. Colored dots use the same color code as in panel (a). Arrows point to representations of *z* = 0 (green) and *z* = *π/*2 (red). The dashed gray curve shows the input manifold projected onto the first two PCA components, which span the same two-dimensional subspace; the two projections may differ by a rotation but both reveal the manifold’s geometry. (d) Square root of the pullback metric, 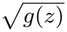, of the input manifold (solid black curve, metric divided by *K/*2) and the neural manifold (gray dashed line) as a function of *z* (strip of colored dots indicates equidistant values). The constant neural metric (line element 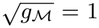) reflects that the feedforward transformation was constructed to isometrically embed the circular stimulus manifold.

with 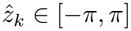 denoting the preferred stimulus of input unit *k*. We will exemplify transformations of representational geometry (i.e., the metric tensor) using a nonuniform distribution of preferred stimuli. We therefore randomly sample 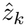 from a von Mises distribution [29] centered at *z* = 0 and with concentration parameter *κ* = 5 (Fig. 2(b) top). This yields a population in which the tuning curve maxima cluster around *z* = 0.

The population activity ***ψ***(*z*) = (*ψ*_1_(*z*), …, *ψ*_*K*_(*z*))^⊤^ forms a (one-dimensional) ring manifold that lies in a two-dimensional linear subspace of ℝ^*K*^ . Indeed, decomposing the cosine tuning curves as

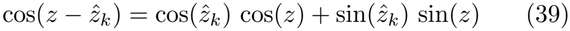

shows that the activity lies in the subspace spanned by the two *K*-dimensional vectors ***ĉ*** and ***ŝ***with entries cos 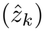 and sin 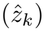, respectively, and that the corresponding “latent variables” *f*_1_(*z*) = cos(*z*) and *f*_2_(*z*) = sin(*z*) form a circle or ring in this space (Fig. 2(a)). To visualize the geometry of this ring in the input space ℝ^*K*^, the geometric arrangement of the vectors ***ĉ*** and ***ŝ***can be taken into account by orthonormalizing ***ĉ*** and ***ŝ***(e.g., via the QR decomposition), resulting in an orthonormal basis for the subspace. The coordinates 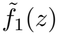 and 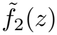 in this orthornomal basis are shown in Fig. 2(c), demonstrating the distortion of the input ring manifold induced by the bias of preferred stimuli.

As a consequence, distances between input activity patterns for neighboring stimuli at 0 and *π* are smaller than those for stimuli at ±*π/*2 (Fig. 2(c)). This is reflected by a non-uniform pullback metric *g*_*J*_ (*z*), Eq. (6), of the input manifold (Fig. 2(d)) with highest resolution around *z* = ± *π/*2 and lowest resolution forstimuli around *z* = 0, where most tuning curves have their peaks. This, at first, counter-intuitive result directly reflects the nature of Fisher information as a measure of stimulus sensitivity [30]: the population activity changes most in the vicinity of ±*π/*2.

In the ring geometry considered so far, the distortion originates from the nonlinearity and bias of the tuning curves. However, geometry can even be reshaped by a purely linear transformation. To show that, we linearly matrix ***ĉ, ŝ***. As a result, the distorted ring in intransform the input activity ***ψ***(*z*) into *N* -dimensional neural activity ***x***^*\**^(*z*) with a (rank-2) feedforward matrix *M* that maps the vectors ***ĉ*** and ***ŝ***to two orthonormal vectors in ℝ^*N*^, using the pseudoinverse of the *K* × 2 put space is transformed into an round circle in neural space, i.e. the unit circle in the subspace spanned by the orthonormal vectors *M* ***ĉ*** and *M* ***ŝ***. Accordingly, the neural tuning curves 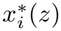 are approximately uniformly distributed across the stimulus range [Δ*π, π*] (Fig. 2(b) bottom). The resulting metric on the ring in neural space is constant (Fig. 2(d)). Hence, all stimuli *z* are equally resolved, assuming, as before, Gaussian noise with stimulus-independent isotropic covariance.

Thus, by linearly mapping low-dimensional nonlinear manifolds *J* ⊂ ℝ^*K*^ to low-dimensional manifolds *ℳ* ⊂ ℝ^*N*^ in the ambient neural space, feedforward synaptic weights can implement nonlinear changes of representational geometry. Beyond reshaping geometry, they can even change the topology of the manifold, as shown for toroidal manifolds in the next subsection.

### C. Toroidal manifolds

#### 1. Representational geometry of grid cell modules

A standard example forming a non-trivial two-dimensional neural manifold are grid cells in the medial entorhinal cortex (MEC) [14, 31–33]. Grid cells exhibit hexagonally arranged firing fields in 2-d arenas (***z*** ∈ℝ^2^) during free foraging behaviors. Grid cells in anatomically neighboring regions of the MEC show the same grid periodicity [34] and are considered to act jointly as functional modules. Here, we follow the general approach [14, 35] and consider the neural manifold generated by a module with fixed grid period.

The activity pattern of one grid cell *i* in a module with period *ℓ* can be phenomenologically described by a superposition of three plane waves [35, 36]:

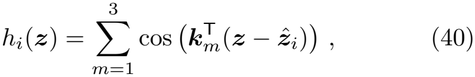

where 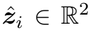 is a cell-specific two-dimensional spatial offset (typically called the grid phase). Such a grid pattern is shown in Fig. 3(a). The grid period and orientation are determined by the wave vectors

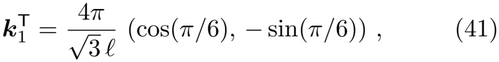

**Figure 3.**
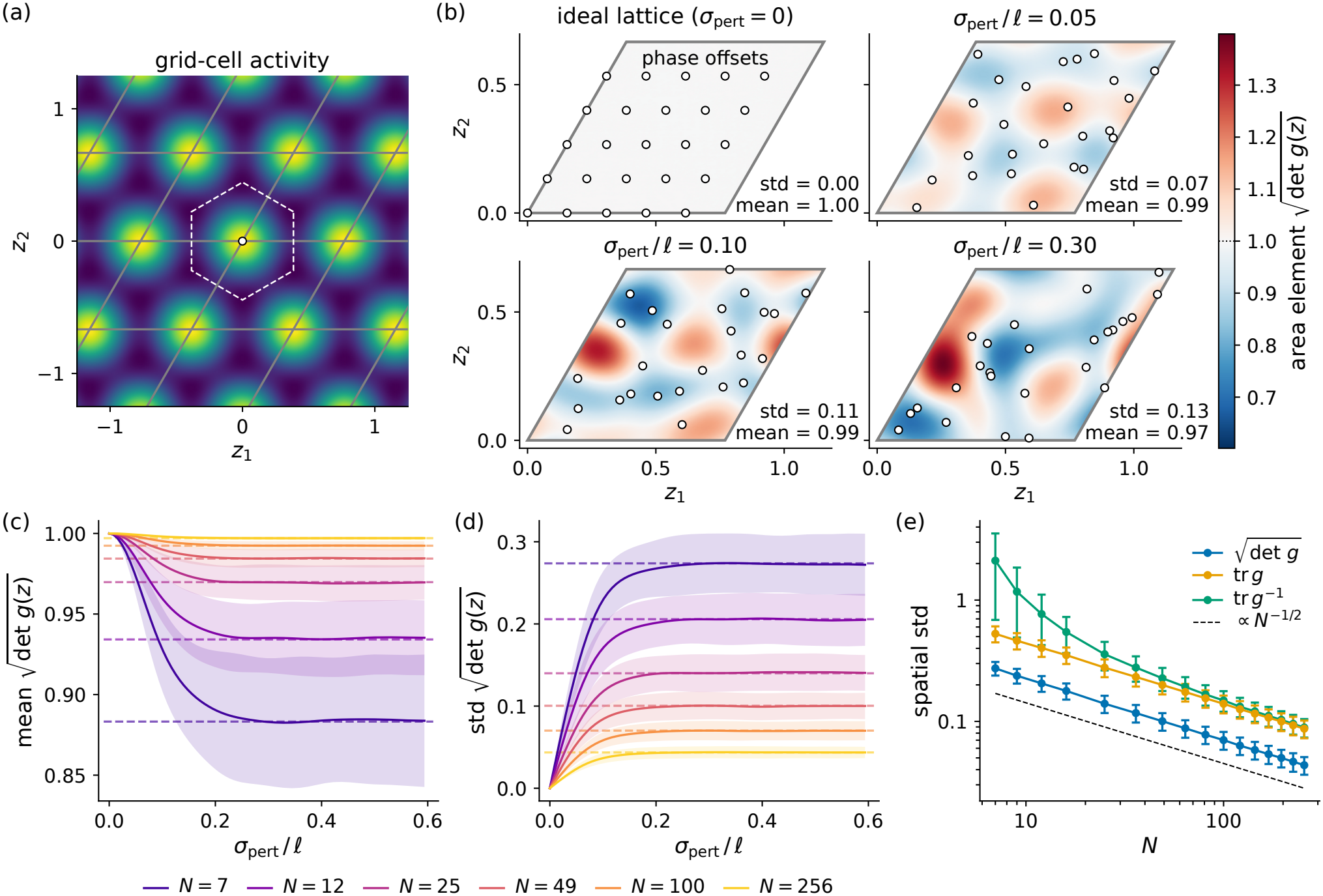
Geometry of gridcell population activity. (a) Grid cell activity modeled as a superposition of three plane waves metric with grid period 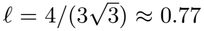, shown here for zero spatial phase offset 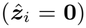. Gray lines indicate the parallelogram tiling of ℝ^2^, and the dashed hexagon outlines a Wigner-Seitz unit cell. (b) Area element 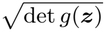 of the normalized tensor for *N* = 25 grid cells with phase offsets arranged on a hexagonal lattice in the parallelogram unit cell, and for isotropic Gaussian perturbations of increasing standard deviations (*σ*_pert_*/ℓ* = 0, 0.05, 0.10, 0.30) applied independently to each offset and mapped back into the unit cell. All subpanels share a common color bar. Inset values give the spatial mean and standard deviation of 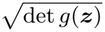over the unit cell. (c) Spatial mean and (d) standard deviation of the area element as a function of perturbation strength for *N* = 7–256 grid cells (colors) per module. Solid curves show the sample mean and shaded bands the sample standard deviation across 1, 000 random realizations of the perturbations. Dashed lines show the sample mean across 1, 000 realizations of *N* grid cells with random phases distributed uniformly in the unit cell. (e) Spatial standard deviations of 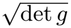, tr *g*, and tr *g*^*−*1^ as a function of *N* for uniformly random phase offsets. All three quantities scale approximately as *N* ^*−*1*/*2^ (dashed reference line). Dots show sample means and error bars show standard deviations across 1, 000 realizations.

***k***_2_ and ***k***_3_, which are rotated from ***k***_1_ by 60^*°*^ and 120^*°*^, respectively. Thegrid period *ℓ* is related to the wave-length as 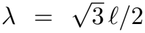. In a single module of grid cells, both grid period and orientation are fixed for all cells in the module. Hence, the population activity ***h***(***z***) = (*h*_1_(***z***), …, *h*_*N*_ (***z***))^⊤^ produces a (at most) two-dimensional neural manifold which depends on the spatial phase offsets 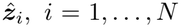. Due to the periodicity of the grid pattern, it suffices to consider the offsets within a (primitive) unit cell of the hexagonal lattice; see Fig. 3(a). In the following, we analyze how the geometry depends on the distribution of the phase offsets in a module.

As before, we analyze the geometry via the pullback metric *g*(***z***) resulting from the Euclidean inner product on the neural activity space ℝ^*N*^, see Eq. (6). If the pull-back metric is constant and proportional to the 2 × 2 identity matrix 1_2_, we consider the neural manifold to be a scaled (local) isometry of the Euclidean plane, faithfully representing distances and angles. Deviations from the scaled isometry property can then be characterized relative to the identity matrix.

A simple way to obtain such a scaled isometry is to choose phase offsets 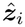 that form a hexagonal lattice and that equally tile a primitive unit cell of the hexagonal grid pattern (Fig. 3(b) upper-left for the parallelogram unit cell). This requires a square number of grid cells, e.g. *N* = 9, 16, 25, By rotating the hexagonal lattice of phase offsets relative to the orientation of the grid pattern, other solutions are also possible; the smallest numbers are *N* = 7 [35] and *N* = 12 (see App. C) . In App. C, we also compute the pullback metric and show the scaled isometry property for the hexagonal lattice configuration, which yields the identity matrix 1_2_ scaled with the prefactor *N* (2*π/ℓ*)^2^ as the metric. In Fig. 3, we therefore normalize the metric by this prefactor.

The homogeneity of the spatial code is illustrated by the area (volume) element 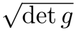 of the normalized metric in Fig. 3(b). Interestingly, a constant and thus flat metric of the neural manifold in the high-dimensional neural activity space (also called conformal isometry between physical and neural space) can be obtained with only a small number of neurons per module; see also [35] and App. C. We test the robustness of this phenomenon by computing the representational geometry under perturbations of these ideal phase configurations. To this end, we jitter each phase offset 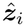 by an independent isotropic Gaussian noise. As we increase the amplitude of the phase jitter, the metric deviates more and more from a scaled isometry (Fig. 3(b)), with smaller resolution, i.e. area element 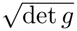, at positions where phase offsets 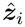 cluster (the peaks of grid fields) and higher resolution at the “population slopes” (cf. Fig. 2).

The spatial mean and standard deviation of the area element depend rather sensitively on the jitter amplitude (Fig. 3(c,d)); they quickly converge to the expected values obtained for randomly distributed phases (dashed lines). However, the deviation from the ideal case decreases with the square root of the number of phases *N* (grid cells per module). Indeed, for the more general case of random phases, the isometry, as assessed by the spatial standard deviation of 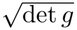, tr *g* and the CramérRao bound tr *g*^*Δ*1^, improves with 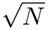 (Fig. 3(e)). We thus conclude that already for *N* = 100 cells per module, the deviations from isometry are sufficiently small (about 7 %) for randomly distributed phases and would not warrant fine-tuned and rather sensitive ideal phase configurations.

While the area element 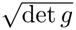 reflects the local expansion of the neural manifold, it depends on how the stimulus space is mapped onto the manifold and is not invariant under a change of coordinates. A coordinate-independent measure of intrinsic geometry is the Ricci scalar curvature *R* [23, 37]), built from up to second derivatives of the metric (App. D. In two dimensions, *R* equals twice the Gaussian curvature, positive in locally sphere-like (elliptic) and negative in saddle-like (hyperbolic) regions, and identically zero for a flat manifold. For a single realization of *N* = 25 grid cells with random phase offsets (Fig. 4(a,b)), *R*(***z***) takes both signs across the unit cell, as it must for a non-flat torus. By the Gauss–Bonnet theorem,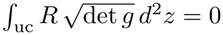.

**Figure 4.**
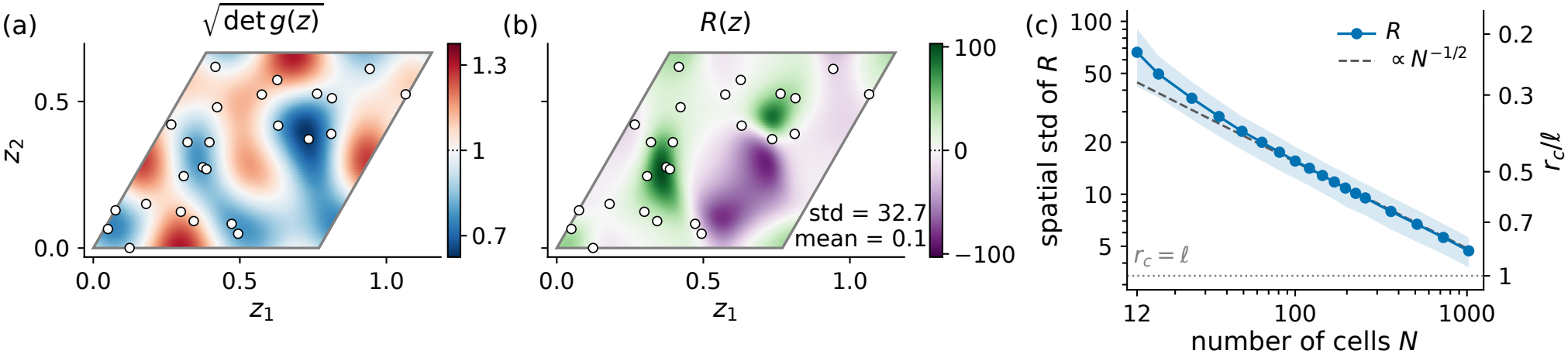
Intrinsic curvature of grid-cell module activity. (a) Area element 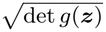 of the normalized metric tensor and (b) the corresponding Ricci scalar curvature *R*(***z***) (App. D) for a single realization of *N* = 25 grid cells with uniformly distributed random phase offsets 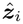 (white circles). The color bar in (a) is centered at 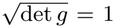 (scaled isometry, cf. Fig. 3(b)). *R* is positive (green) in locally sphere-like and negative (violet) in locally saddle-like regions; it is largest where phase offsets cluster and shows no apparent relation to the area element. Inset: spatial standard deviation and mean of *R* over the unit cell. As required for a torus, 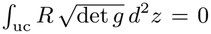 (Gauss–Bonnet; numerical error *<* 10^*Δ*14^). (c) Spatial standard deviation of *R* as a function of *N* : mean (markers) and standard deviation (shaded band) across 1, 000 realizations. Dashed line: ∝ *N* ^*Δ*1*/*2^, approached for large *N* . Right axis: typical curvature radius *r*_*c*_ = 2*/* std *R* in units of the grid period *ℓ*; dotted line, *r*_*c*_ = *ℓ*.

The curvature shows no apparent relation to the area element (Fig. 4(a,b)). Instead, *R* tends to be positive where phase offsets cluster, and the manifold bulges there like a sphere. In between phase offset clusters, *R* tends to be negative. The spatial standard deviation of 32.7 corresponds to a typical local radius of curvature 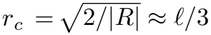.

As for the area element, the spatial standard deviation of *R* decreases as *N*^*Δ*1*/*2^ for large *N* (Fig. 4(c)), so the curvature radius grows only as *r*_*c*_ ∝ *N* ^1*/*4^. Unlike the area element, however, whose deviation from isometry is only ≈ 2% at *N* = 1024, the curvature stays appreciable, with *r*_*c*_ ≈ 0.85 *ℓ < ℓ*: as a second derivative of the metric, *R* is of order |***k***_*m*_|^2^ times the relative metric deviation, and since 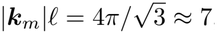, this is large on the scale of the unit cell even when the deviation is small. The manifold is thus metrically close to a scaled isometry yet intrinsically curved on the scale of its own size. Hence, already in the phenomenological grid-cell model, random phase offsets yield a manifold whose geometry differs from that of the Euclidean plane.

#### 2. Feedforward model of grid cells

Although the phenomenological grid-cell activity model based on Eq. (40) already allows us to study its representational geometry, it does not provide mechanistic insights into contributions from synaptic weight matrices. As a first approach to include network mechanisms, we consider a purely feedforward model. In the case of propagated slow noise, the recurrent weights do not change the Fisher information (App. B) and they can thus be set to zero (for the effect of recurrence under fast noise, see Sec. III D). Previous feedforward models have constructed grid cells from place cell inputs [38–42]. However, place-field-like inputs are not necessary to generate grid cells [43]. Here, we show that already a fixed Hebbian feedforward transformation of random spatial inputs suffices, as long as the inputs ***ψ***(***z***) contain suitable spatial information.

Input tuning functions

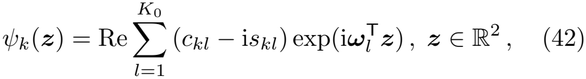

are modeled as stationary random fields constructed from *K* = 2*K*_0_ random Fourier features [44] with *K*_0_ wave vectors ***ω***_*l*_ drawn independently from an isotropic two-dimensional Gaussian distribution with mean zero and standard deviation *Σ*_*ω*_ (spatial frequency scale) and weights *c*_*kl*_, *s*_*kl*_ obtained as the elements of a *K* × *K* random orthogonal matrix. Since the inputs should contain spatial frequencies near that of the grid cells, we set 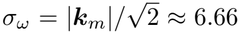, the value for which the Gaussian density of ***ω***_*l*_ at the grid wave vectors ***k***_*m*_, Eq. (41), is maximized. Examples are shown in Fig. 5(a).

**Figure 5.**
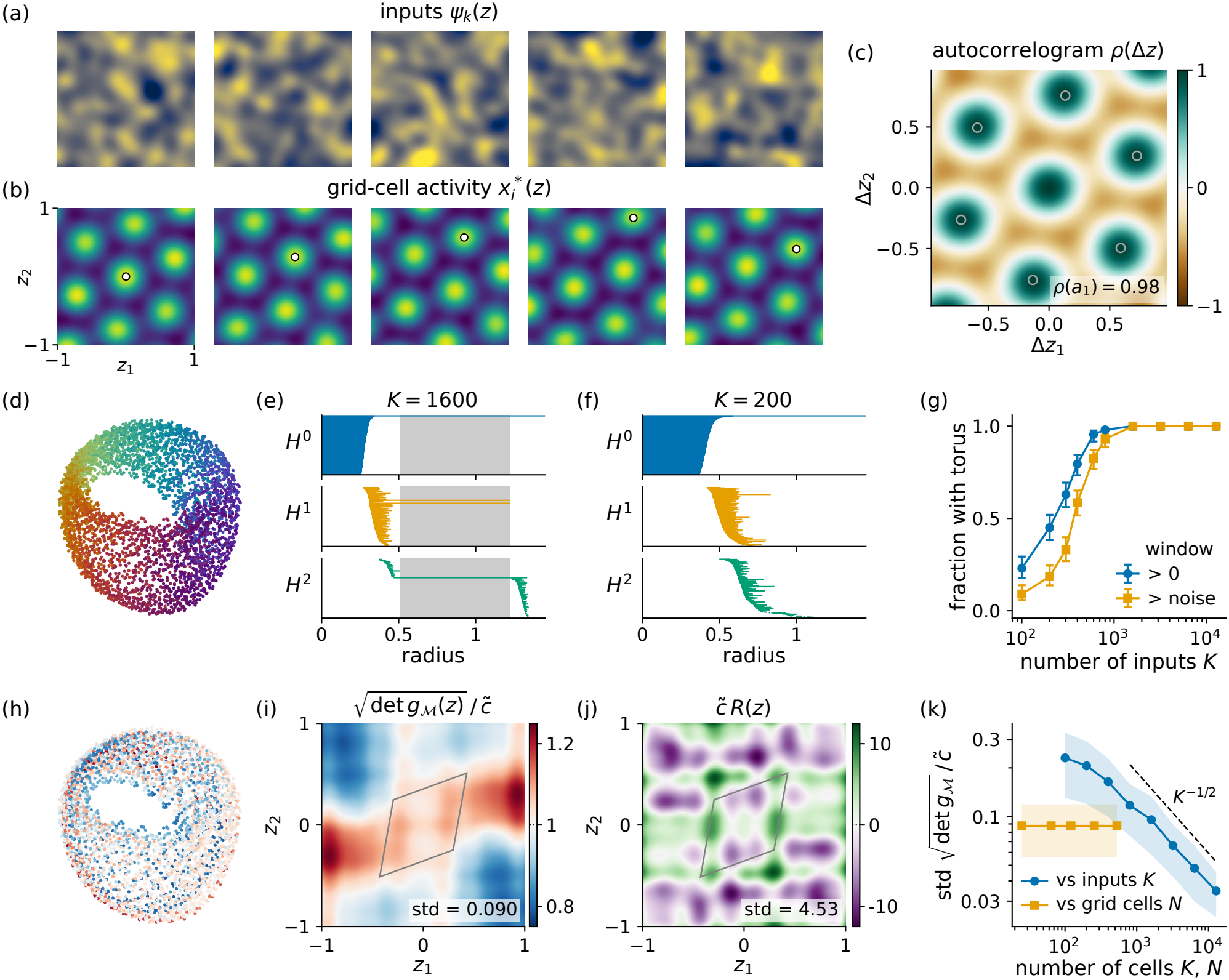
Feedforward model of a grid-cell module: activity, topology, and geometry. Results are obtained for *N* = 256 target grid patterns *h*_*i*_(***z***) with period 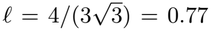, from Eq. (40), and grid orientation 20^*°*^. The 16 × 16 phase offsets 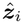 are regularly spaced in the parallelogram unit cell. As an input population we use *K* = 1600 random spatially modulated inputs *ψ*_*k*_(***z***), Eq. (42). Panels (a)–(f), (h)–(j) are obtained from one realization, on the analysis box [− 1, 1]^2^; (g),(k) are obtained from ensembles of 200 realizations per point. (a) Five exemplary input tuning curves *ψ*_*k*_(***z***) with random spatial modulations. (b) Five exemplary grid-cell activities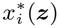; white dot indicates target phase offset 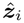. (c) Population activity autocorrelogram *ρ*(Δ***z***). Averages are taken over the stimuli with ***z*** and ***z*** +Δ***z*** both in the box. Circles: the six target lattice vectors ***a***_1_, ± ***a***_2_, ± ***a***_3_, of length *ℓ* and at 20^*°*^, 80^*°*^, and 140^*°*^. (d) Isomap embedding of the population activity into 3-d at 4225 stimulus positions (6 principal components, 10 neighbors), colored by the grid phase 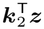 wrapped to [− *π, π*). (e),(f) Persistence barcodes (Vietoris–Rips, ℤ_3_ coefficients, 450 points subsampled from the 4225): lifetime of components (*H*^0^), loops (*H*^1^), and cavities (*H*^2^) against the filtration radius, in units of the rms activity; a torus gives one, two, and one long bar. Bars are sorted within each strip, so their vertical position carries no meaning. Shaded: the radii with the exact 1, 2, 1 configuration. (e) Example realization with *K* = 1600. (f) same as (e) for *K* = 200 (median realization). (g) Fraction of 200 realizations whose barcode shows a torus: at some radius (window *>* 0), or also outlasting the longest non-torus bar in *H*^1^ and *H*^2^ (window*>* noise). Wilson 95 % intervals [45]. (h) The embedding of (d), recolored by the area element of (i) and sharing its color bar. (i) Area element 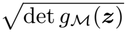, normalized by 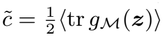 over the box; gray: a unit cell of the grid lattice; annotation: the spatial standard deviation. (j) Ricci scalar curvature *R*(***z***) of the same metric, multiplied by 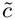; same unit cell and annotation. (k) Spatial standard deviation of the area element against the number of inputs *K* (at *N* = 256) and of grid cells *N* (at *K* = 1600); markers, sample means; bands, sample standard deviations over 200 realizations; dashed, the *K*^*−*1*/*2^ slope.

We define a Hebbian feedforward weight matrix

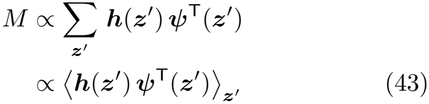

as a sum of outer products of the spatially modulated input patterns ***ψ***(***z***^*′*^) and grid patterns ***h***(***z***^*′*^) given by Eq. (40) with grid period 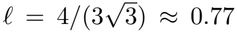 (wavelength *λ* = 2*/*3). For this construction, stimuli (locations) ***z***^*′*^ are uniformly distributed on a square training box [Δ1.3, 1.3]^2^ ⊂ℝ^2^, which is padded by two input correlation lengths, 2*/Σ*_*ω*_, around the analysis box [Δ 1, 1]^2^.

For numerical calculations, we sampled ***z***^*′*^ from a square grid with high resolution. The *N* = 256 target grid patterns *h*_*i*_(***z***^*′*^) are obtained using equally spaced phase offsets 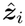 in the unit cell that would result in an ideal scaled isometry, as discussed above. With ***x***^*\**^(***z***) = *M* ***ψ***(***z***), the random spatial code is transformed into a hexagonal gridcell code with the target grid period *ℓ* (Fig. 5(b)).

The autocorrelogram *ρ*(Δ***z***) of the grid-cell module activity shows clear hexagonal periodicity (Fig. 5(c)), indicating a toroidal neural manifold. Isomap embedding [46] into 3-d indicates a torus with points organized by a grid phase (Fig. 5(d)).

We corroborate the toroidal topology with persistent (co-)homology analysis [47, 48]. The persistence barcodes for *K* = 1600 input cells show a long window of radii in which the 1, 2, 1 configuration of a torus is expressed (Fig. 5(e)). This means that the neural manifold has toroidal topology over a large range of scales. However, the toroidal topology breaks if the number of inputs *K* is too low, as exemplified for *K* = 200 in Fig. 5(f). In ensembles of 200 realizations of the *K* input neurons, we quantify the fraction with a torus configuration present in the barcode. As a criterion, we consider both a window of radii of any length larger than zero (blue curve in Fig. 5(g)) and a robust window whose length must be longer than the remaining non-torus (noise) bars (orange curve in Fig. 5(g)). With *K*≥ 1600 input neurons, the persistence barcodes robustly indicate a torus (Fig. 5(g)). Notably, the toroidal structure is induced solely by the symmetry of the target grid patterns ***h***(***z***^*′*^) in the feedforward weights Eq. (43), without requiring recurrent connectivity or a continuous attractor network.

While it is well-established [14, 35] that the toroidal topology of grid-cell module activity arises from their hexagonal symmetry, little attention has been paid so far to the topology of the geometric deviations from a scaled isometry (but see [49]). On a true torus (as, e.g., generated by a perfect continuous attractor manifold), also the deviations must be periodic. For a feedforward model as constructed here, however, the toroidal topology is induced by a linear transformation of the inputs with non-periodic tuning curves *ψ*_*k*_(***z***), and thus deviations do not need to follow the torus. We illustrate this by coloring the Isomap embedding according to the area element of the pullback metric of the feedforward model (Fig. 5(h)), which reveals that the manifold consists of layers wrapped around a torus rather than being a plain torus. The layers reflect that the area element is not a function on the torus but a function on the plane, here shown on the analysis box [− 1, 1]^2^ (Fig. 5(i)). The same holds for the Ricci scalar curvature (Fig. 5(j)).

We note that the inversion symmetry of both area element and scalar curvature (Fig. 5(i,j)) arises from the construction of the random fields in Eq. (42) together with the centered square training box [−1.3, 1.3]^2^ used for the feedforward matrix *M* .

Lastly, we ask how these deviations depend on the number of inputs *K* and the number of grid cells *N* (Fig. 5(k)). The number of grid cells *N* has no influence (Fig. 5(k)): all grid cells combine the same three plane waves as estimated from the inputs, so the estimation errors are shared by all cells and do not average out. For the lattice phase offsets used here, the combination itself would result in an exact scaled isometry (Sec. III C 1), so that *N* only rescales the metric and the inhomogeneity stems entirely from the feedforward transformation. The plane waves are estimated from *K* random features, and for large *K*, the spatial standard deviation of the area element decreases as *K*^*−*1*/*2^. This reflects the outer-product construction, Eq. (43), which ignores the correlations between the inputs, and mirrors the *N*^*−*1*/*2^ scaling for random phase offsets (Fig. 3(e)). With *K* = 1600 input neurons, the deviations from isometry are still about 10 %. These deviations could thus be observable in experimental data; since they are not functions on the torus, they would be tied to position rather than to grid phase.

### D. Fast noise and recurrent connectivity

So far, we only considered slow input noise with constant covariance ∑^*′*^(*t*) = ∑^*′*^ and concluded that in this case, the recurrent connectivity does not affect the statistical manifold. However, this no longer holds for fast noise. As an illustration, we assume Gaussian white noise that is isotropic in neuron space:

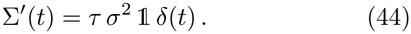

Following the derivation in App. B, we obtain the term in the Fisher information that depends on the recurrent weight matrix *W*, Eq. (B8), as

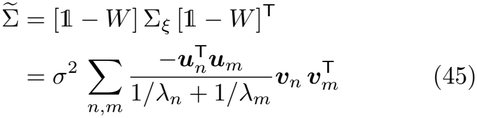

with the eigenvalues *λ*_*n*_ and the corresponding right- and left-eigenvectors ***v***_*n*_ and ***u***_*n*_ of the matrix *W*− 1. Therefore, the Fisher information for propagated fast noise, Eq. (32), reads

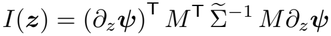

and depends on the eigenvectors and eigenvalues of *W* . We will explore this *W* -dependence in two examples.

#### 1. Rank-1 recurrent weights

In the simple case of recurrent weights *W* = ***w w***^T^ with ***w*** ∈ℝ^*N*^, the matrix *W*− 1 has the eigenvalue *λ*_1_ = |***w***| ^2^ −1 with corresponding eigenvector ***w***, spanning a one-dimensional subspace. All other (orthogonal) subspaces have the eigenvalues *λ*_*n*_ = −1, *n* ≥2. Since *W* − 1 is symmetric, we can choose an orthonormal basis of eigenvectors ***v***_*n*_ starting from ***v***_1_ = ***w****/* |***w***| . Then, ***u***_*n*_ = ***v***_*n*_ and Eq. (45) becomes

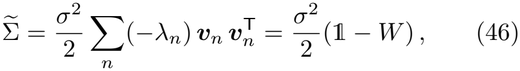

which in fact, with isotropic white noise, holds for any symmetric *W* .

With Eq. (46), we obtain the Fisher information

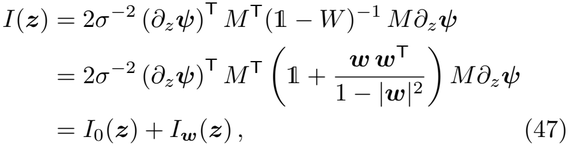

where *I*_0_(***z***) is the (fast-noise) Fisher information without recurrent weights, i.e. for ***w*** = 0, and

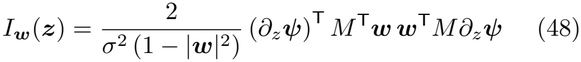

can be interpreted as the Fisher information available after projecting the local signal direction *M∂*_*z*_*ψ*(***z***) on the subspace spanned by ***w***. Since |***w***| ^2^ *<* 1 to ensure stability of the fixed point, both contributions are non-negative, and for ***w*** with ***w***^T^*M* ≠ **0**, the full Fisher information (47) can be larger than *I*_0_(***z***). Specifically, regions of the feedforward manifold *Mψ*(***z***) where *M∂*_*z*_*ψ*(***z***) is aligned with ***w*** will exhibit improved resolution and statistical discriminability; see also [50].

A similar effect will be illustrated next for more complex symmetric matrices.

#### 2. Grid cells with recurrent connectivity

As a second example for the impact of recurrent connectivity under fast noise, we revisit the grid-cell network introduced in Sec. III C 2. We introduce a recurrent weight matrix *W* that is constructed through a Hebbian outer-product rule

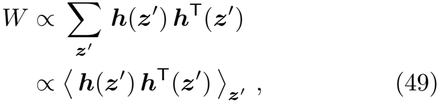

based on the (target) grid patterns ***h***(***z***) = (*h*_1_(***z***), …, *h*_*N*_ (***z***))^T^, ***z*** ∈ ℝ^2^, of the cells, which are defined by phase offsets 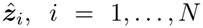 . The angular brackets in Eq. (49) denote a spatial average over a set of points ***z***^*′*^∈ ℝ^2^. This construction ensures that all dimensions spanned by the feedforward input *M* ***ψ***(***z***) also affect the recurrent dynamics. To focus on the impact of the recurrent connectivity, we simplify the feedforward input by using the grid patterns ***h***(***z***) themselves, i.e., without recurrent weights, the steady state population activity is ***x***^*\**^(***z***) = ***h***(***z***), which we analyzed in Sec. III C 1.

If we take the spatial average over the uniform distribution on a (parallelogram) unit cell of the hexagonal grid pattern (Sec. III C 1), the resulting correlation matrix has the form

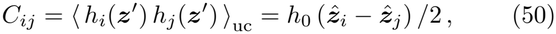

where 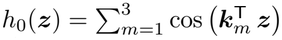is the grid pattern from Eq. (40) with zero phase offset. The recurrent weight matrix *W* can therefore be written as

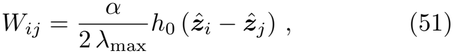

where the factor *α/λ*_max_ is chosen such that the largest eigenvalue of *W* is *α* ∈[0, 1), guaranteeing stable fixedpoint dynamics. While being constructed through an outer-product rule, Eq. (49), the weight matrix *W* exhibits spatial structure in the space of the grid phase off-sets (Fig. 6(a)). Indeed, according to Eq. (51), the weight matrix *W* is proportional to the grid pattern *h*_0_(***z***) itself, evaluated at the phase offset differences 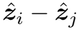 between the neurons. This gives rise to translation-invariant connectivity: the weights depend only on the phase-offset difference 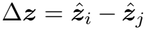, and the corresponding weight kernel in the unit cell of the hexagonal grid is of “Mexicanhat” type (Fig. 6(a), right), as in continuous attractor network models [51, 52].

**Figure 6.**
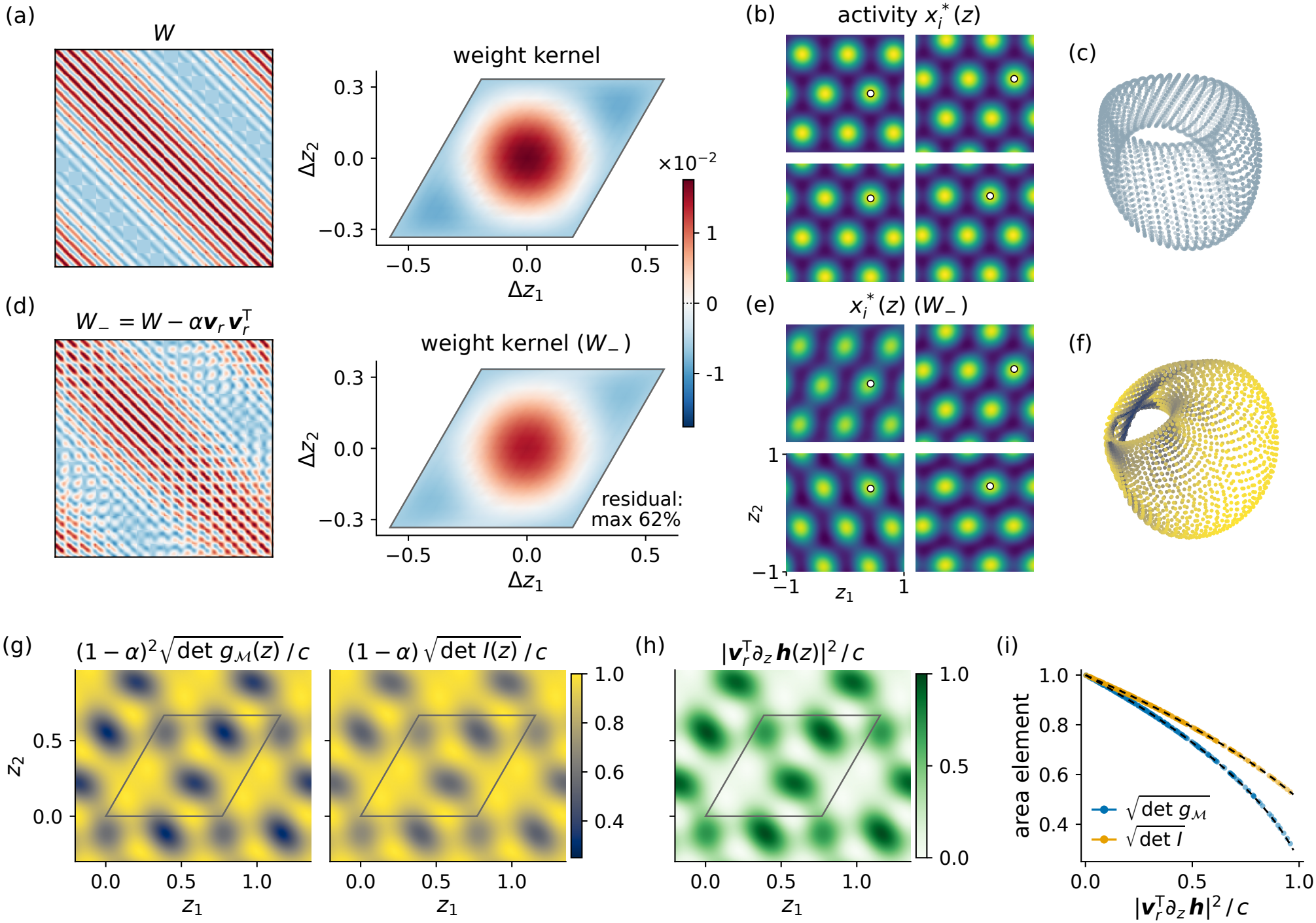
Representational geometry of a grid-cell module with recurrent connectivity. *N* = 256 cells with phase offsets 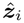 on a hexagonal lattice and recurrent weights 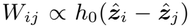, Eq. (51), with largest eigenvalue *α* = 0.75. (a) Left: recurrent weight matrix *W* (entry (1, 1) top left). Right: weight kernel, the mean of *W*_*ij*_ over all neuron pairs with a given phase-offset difference 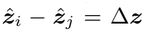 in the unit cell. For *W*, every such pair has the same weight, Eq. (51), so the “Mexican-hat” type kernel is exact. All four maps in (a),(d) share one color bar. (b) Steady-state activity 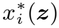 of four cells; white dot: phase offset 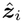. Shared color scale (dark blue to yellow) with (e). (c) Isomap embedding of the neural manifold into 3-d (*k* = 6 nearest neighbors); uniform color, as *W* acts as a scaled identity on the span of ***h***(***z***) and the area element is constant. (d)–(f) As (a)–(c), for 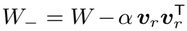 with a random unit vector ***v***_*r*_ in the eigenspace of the 6-fold degenerate eigenvalue *α*. The weight kernel keeps its shape but is scaled by 5*/*6, while individual entries (*W*_*−*_)_*ij*_ deviate from it by up to 62 % of the *W* -kernel peak (annotation), quantifying the broken translation invariance. The torus in (f) is colored by the area element. (g) Area element of the neural, 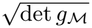, and of the statistical manifold, 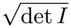, for *W*_*−*_, each normalized by its constant full-*W* value, (1 − *α*)^*−*2^ *c* and (1 − *α*)^*−*1^ *c* with the isometry prefactor *c* = *N* (2*π/l*)^2^ (Sec. III C 1); the reference is therefore 1. (h) Normalized squared overlap 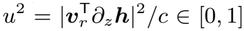 of the local signal directions with the removed eigenvector. Dark color marks a strong effect of the removal in both (g) and (h). (i) Normalized area elements from (g) versus *u*^2^ from (h): markers, numerical values from the unit cell; dashed, the exact relations 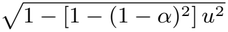 (neural) and 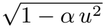 (statistical). The smaller coefficient, *α* instead of 1 − (1 − *α*)^2^, is the attenuation by the Fisher information.

With recurrent weights, the steady state population activity and the neural manifold are then given by ***x***^*\**^(***z***) = (1 − *W*)^*−*1^***h***(***z***). The individual neurons show grid-cell activity patterns (Fig. 6(b)) and the manifold has a toroidal shape (Fig. 6(c)). Since the square matrix (1− *W*)^*−*1^ is invertible, it cannot change the topology.

The symmetric matrix *W* given by Eq. (51) has in general rank 6 [e.g., see Eq. (40)] and the spectral decomposition

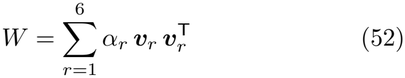

with the nonzero, possibly degenerate, eigenvalues *α*_*r*_ and corresponding orthonormal eigenvectors ***v***_*r*_. The outer-product rule, Eq. (49), implies that these eigenvectors also yield an orthonormal basis of the 6-dimensional subspace spanned by the input grid patterns ***h***(***z***). For hexagonal lattice phase offsets 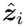 (see Sec. III C 1), the largest eigenvalue *α* is 6-fold degenerate and 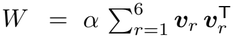 is a scaled orthogonal projection. In this case, illustrated in Fig. 6(a–c), the neural manifold is just the scaled feedforward input: ***x***^*\**^(***z***) = (1 −*W*)^*−*1^***h***(***z***) = (1 − *α*)^*−*1^***h***(***z***) with the gain factor (1 − *α*)^*−*1^ *>* 1, demonstrating how recurrent feedback can increase the size of the manifold.

For isotropic Gaussian white noise with covariance (44), the Fisher information of our grid-cell network with recurrent connectivity is given by

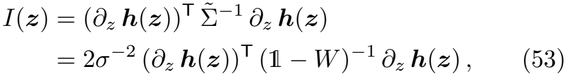

where we used that the effective covariance matrix 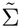 from Eq. (45) reduces to 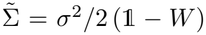 for general symmetric *W*, see Eq. (46). We set *σ*^2^ = 2 in the following.

A statistical manifold ***y***(***z***) that has the Fisher information (53) as its pullback metric can be obtained as

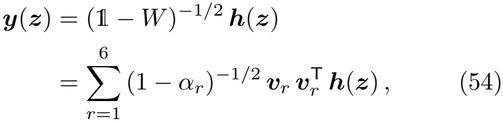

using the spectral decomposition (52) of *W* and that the input manifold ***g****c*(***z***) lies in the span of the eigenvectors ***v***_*r*_, *r* = 1, …, 6 . As mentioned above, for hexagonal lattice phase offsets, the largest eigenvalue *α* is 6-fold degenerate. In this case, the statistical manifold (54) simplifies to ***y***(***z***) = (1 − *α*)^*−*1*/*2^***h***(***z***), the feedforward input scaled by another gain factor. Hence, if the recurrent matrix *W* acts as a scaled identity on the subspace spanned by the feedforward input ***h***(***z***), both the neural and the fastnoise statistical manifold are just scaled versions of the input manifold (Fig. 6(c)).

To further explore the effect of recurrent connectivity on the representational geometry, we manipulate the weight matrix *W* by removing one of the eigenmodes in the spectral decomposition (52), where we set one eigenvalue *α*_*r*_ = 0. Focusing again on the case of a 6-fold degenerate eigenvalue *α*, we first construct a random orthonormal basis of eigenvectors ***v***_*r*_, *r* = 1, …, 6, so that the removed mode corresponds to a random unit vector in the 6-dimensional subspace spanned by the input grid patterns ***h***(***z***). The resulting recurrent weight matrix *W*_*−*_ is visibly distorted relative to *W* (Fig. 6(d), left) while its translation-averaged weight kernel is only reduced in amplitude by a factor of 5*/*6 with an essentially unchanged shape (Fig. 6(d), right). Individual weights (*W*_*−*_)_*ij*_, however, deviate strongly from this averaged kernel: unlike *W*, the matrix *W*_*−*_ is no longer a function of the phase offset difference 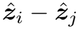 alone.

With *W*_*−*_ as recurrent connectivity, the steady-state activity ***x***^*\**^(***z***) = (1 − *W*_*−*_)^*−*1^***h***(***z***) still resembles gridcell patterns (Fig. 6(e)), which can be partly attributed to the unaltered feedforward input. With Isomap embedding, the toroidal shape of the neural manifold appears distorted (Fig. 6(f)), even though the invertible matrix (1 − *W*_*−*_)^*−*1^ preserves the toroidal topology of ***h***(***z***).

To quantify the distortion of the representational geometry and compare the statistical manifold ***y***(***z***), Eq. (54) with *W*_*−*_, to the neural manifold, we consider the area elements. Removing a mode from the recurrent weight matrix reduces the local area elements 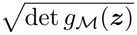 and 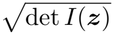 of the neural and statistical manifold, respectively, in certain regions (Fig. 6(g)). As above, we take the full *W* for hexagonal lattice phase offsets as the reference, which yields via ***x***^*\**^(***z***) = (1 − *α*)^*−*1^***h***(***z***) and ***y***(***z***) = (1 − *α*)^*−*1*/*2^***h***(***z***) the constant metric and area element of the scaled isometry ***h***(***z***) from Sec. III C 1, only multiplied by the respective gain factors (1 − *α*)^*−*2^ and (1 − *α*)^*−*1^. After dividing by these gain factors and by the isometry prefactor *c* = *N* (2*π/l*)^2^, the full-*W* reference is 1 in Fig. 6(g). Whereas the spatial patterns of the reduction are identical, the relative decrease of 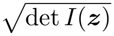 is weaker than that of 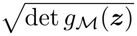 (Fig. 6(g)), arguing that the geometric effect of the mode emoval is attenuated when using the Fisher information (53) as the metric.

To understand how the removed mode is related to the change in geometry, we project the Jacobian (local signal directions) of the feedforward input, *∂*_*z*_ ***h***(***z***) ∈ ℝ^*N ×*2^, onto the removed mode’s eigenvector ***v***_*r*_ (Fig. 6(h)). In regions of the manifold where this overlap 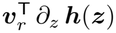 has a large amplitude, the mode contributes the most to the geometry and removing it should have the strongest impact. In fact, the squared norm of this overlap is exactly related to the reduction of local area (Fig. 6(i)). This is analogous to the previous symmetric rank-1 example and Ref. [50], only that here the Fisher information decreases in those regions where the removed mode would otherwise have averaged out the fast noise.

More generally, for isotropic white noise and with symmetric recurrent weights, the Fisher information behaves quite similarly to the pullback metric of the neural manifold. In both cases, the recurrent connectivity amplifies the feedforward input in the directions of its eigenvectors with positive eigenvalues, only with different exponents in the gain factors (see Eq. (54)), thereby increasing the area and resolution of the feedforward manifold.

## IV. DISCUSSION

Representational geometry of neural manifolds is paramount to the study of neural codes in high-dimensional population activity (e.g., [53]), and it is tightly connected to information geometry [18, 19] as already pointed out previously [15]. In this paper, we further elaborated this link for general noise distributions, specifically showing that for stimulus-independent isotropic Gaussian noise, the Fisher information matrix corresponds to the pullback metric of the neural manifold up to a constant factor. More generally, we showed that the Fisher information can be considered as a pullback metric induced by a certain metric on the neural manifold. Under integrability conditions, the neural manifold can be transformed so that the Fisher information is the pullback of the standard Euclidean metric, yielding an embedding of the statistical manifold in the same embedding space as the neural manifold. For rate-based neural networks, we have derived how the metric and Fisher information depend on neural circuit elements such as tuning curves, feedforward, and recurrent synaptic weight matrices, and thereby found that the recurrent weight matrix does not directly affect information geometry for slowly varying noise (e.g., excitability modulations, trial-to-trial variability), whereas it has the beneficial effect of averaging out fast (e.g. white-)noise fluctuations. Beck *et al*. [20] obtained similar results for spiking networks, showing that the covariance structure of the noise determines the effect of recurrent connectivity on Fisher information.

### Feedforward-induced geometry and topology

The independence of information geometry on the recurrent weights for slow noise further emphasizes the importance of geometrical transformations induced by feedforward synaptic weights. Feedforward weights allow transformations of the metric (i.e. manipulations of the resolution) and even the topology of the input manifold, as shown by feedforward models of grid cells where a toroidal topology is created from a finite 2-d square. Despite having toroidal topology, however, grid cell modules can have a flat metric (establishing a conformal isometry [35]) for certain regularly spaced grid phases. Deviations from isometry (and intrinsic curvature) are generated by imperfections, e.g. clustered phases [54] or outer-product feedforward weights derived from random spatial inputs (Fig. 5).

### Population slopes

Our application examples also illustrate how information geometry emphasizes the “slopes of the population tuning”, i.e., stimulus regions and directions with large overall changes of the neural activity, in contrast to the prevailing view that the density of tuning curve peaks reflect the importance of a stimulus feature [55–57].

### Intrinsic vs. extrinsic geometry

The focus on intrinsic geometry takes the perspective of an observer who measures distances along the manifold itself. While this is the natural viewpoint of information geometry [18], it can be challenged by a simple linear readout. A perceptron computes an inner product in the neural embedding space (or at least in the linear subspace defined by the connectome) and is therefore sensitive to extrinsic geometry. For example, on a highly extrinsically curved surface such as a swiss roll, two points may be close in the embedding space but relatively remote on the manifold. A linear readout as in a perceptron cannot extract the intrinsic structure of this curved manifold and instead operates on the geometry in the embedding space [58]. For shallow networks with a linear readout, the focus on intrinsic geometry is therefore not directly justified. Deep neural networks, however, can learn to progressively unfold and flatten nonlinear manifolds, so that intrinsic structure can become accessible to a final linear readout layer [59]. Intrinsic information geometry may thus be available to a deep readout layer as its input may represent the latent variables in a flat Euclidean space.

#### Nonlinear networks and attractors

Throughout, we restricted our analysis to linear network dynamics, omitting a nonlinear activation function *ϕ*(*x*). This corresponds to analyzing manifolds in the vicinity of an attractor state of a nonlinear network, where the dynamics can be approximated by linearization with effective recurrent connections 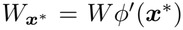 . Such linearization is justified when inputs change slowly with limited dynamic range. However, the linear dynamics is unable to capture the role of unstable fixed points and separatrices, which are usually employed in decision making dynamics, where trajectories are channeled toward one of several stable attractor states. Our results nevertheless have important implications for decision making if we assume such non-linear dynamics in a downstream network. This readout layer will be particularly sensitive to stimulus changes in regions of high Fisher information or high volume element 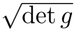, mapping small stimulus differences to large distances in neural activity space.

#### Continuous attractors

In cases of continuous attractors, the formalism can be generalized by having the attractor state ***x***^*^(***z, o***) not only characterized by the stimulus ***z*** but also a parameter ***o*** along the attractor manifold. At each point of the attractor (identified by ***o***) we can again compute the linearized effective recurrent matrix 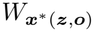 and compute the pullback metric with respect to ***z*** on the subspace orthogonal to the tangent space spanned by 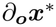 (where 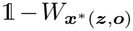 has zero eigenvalue). At any point of the attractor, the stimulus ***z*** may therefore have distinct representational geometry unless the feedforward input *M* ***ψ*** is exactly parallel to the attractor manifold as, e.g., it is assumed in the ring attractor of the head direction system. In this latter case, or more generally, if changes in ***z*** evoke tangent vectors parallel to the attractor manifold, a metric can still be constructed that adds effects of responses orthogonal and parallel to the attractor manifold and contains mixed terms. Thus, even for continuous attractors, information geometry can evaluate the encoding of stimulus space (see also our first analysis of the grid cell metric, where we did not make any mechanistic assumptions); however, the specific relation between metric and circuit parameters *W, M, ϕ, ψ* is less straightforward and will generally require additional assumptions on how changes in the input affect the position on the attractor manifold, as, e.g., how changes in position move bumps on the ring attractor [60, 61].

#### Outer-product networks

In the example applications, we have focused on synaptic weight matrices of outerproduct form, in which the number of linearly independent vectors is typically much smaller than the number of neurons. Such low-rank networks have recently gained considerable interest as they provide a mechanistic account of low-dimensional manifolds [62, 63]. Our main motivation to use outer-product matrices was, however, to better be able to interpret and visualize the results. Outer-product matrices are not necessarily low-rank (if the number of linearly independent summands is of the same order as the number of neurons), but are often considered to be biologically plausible as they can be explained as resulting from local Hebbian learning rules [64]. Our general results on information geometry never made use of the low-rank property and thus also apply, e.g., to sequence generating networks with as many outer product summands as sequence transitions [65–68].

#### Outlook: Dynamic attractors

Sequence-generating networks are examples of networks that not yield an attracting fixed-point state ***x***^*^(***z***) but rather generate attracting trajectories [69, 70]. Information geometry of such dynamical manifolds needs to consider time as another intrinsic dimension [71]. A generalization of our framework to such temporally evolving manifolds will be an important next step, enabling us to also treat the geometry of neural manifolds that evolve along motor trajectories [72], or more generally, time-evolving stimuli.

## ACKNOWLEDGMENTS

This work has been funded by the German Research Association (DFG) under grant numbers LE2250/23-1 and Inst. 514483642 (CRC TRR 384/1 2024 IN-CODE).

## Appendix A: Derivation of the effective noise covariance Q

The solution of the noise-driven linear dynamical system (Eq. (28))

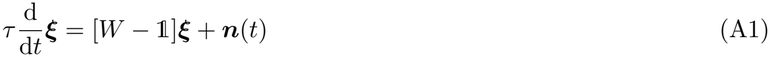

is obtained in closed form via the matrix Green’s function exp([*W* − 1]*t/τ*) as

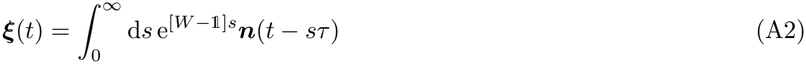

with scaled time *s* = *t/τ* . We therefore compute the covariance between network response ***ξ*** and noise ***n*** as

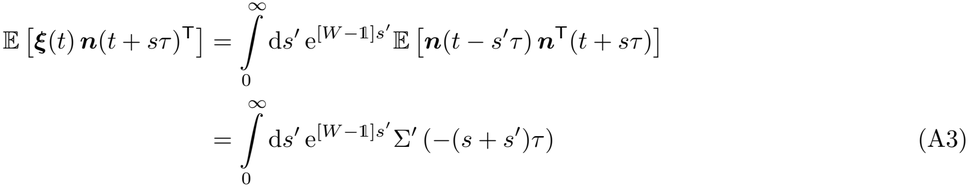

with the covariance matrix Σ^*′*^ of ***n***(*t*). For *s* = 0 and with the symmetry E [***ξ***(*t*) ***n***(*t* + *sτ*)^T^]^T^ = E [***n***(*t*)***ξ***(*t − sτ*)^T^], we arrive at the expression for the effective noise covariance *Q* in Eq. (30).

## Appendix B: Fisher information under recurrent propagation of noise

### Spectral decomposition

If the matrix *A* = *W* −1 is diagonalizable with eigenvalues *λ*_*n*_, *n* = 1, …, *N*, we can write its spectral decomposition as

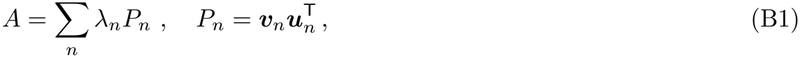

with the (right-)eigenvector ***v***_*n*_ and the corresponding dual basis vector ***u***_*n*_, i.e., 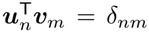 and thus *P*_*n*_ *P*_*m*_ = *δ*_*nm*_ *P*_*n*_. Furthermore, Σ_*n*_ *P*_*n*_ = 1, since the eigenvectors form a basis. The eigenvectors of *A* are the eigenvectors of the recurrent weight matrix *W* . If *W* = 0, i.e. in the absence of recurrent connections, the eigenvalues are *λ*_*n*_ = −1 for all *n* and we can choose any basis of vectors in the following.

Using this decomposition, we can express the noise covariance Σ^*′*^(*t*) in Eq. (30) in terms of the eigenvectors of *W* as

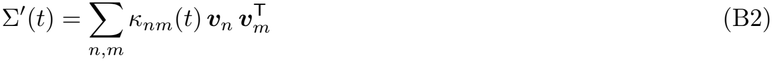

where 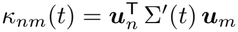. With this expression, we obtain the effective noise covariance, Eq. (30), as

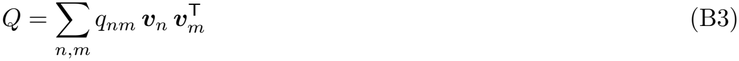

with the coefficients (matrix elements)

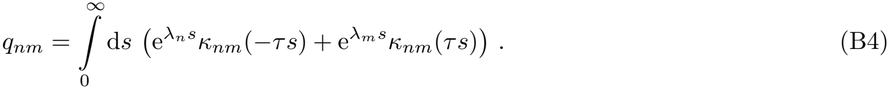

Note that for temporally correlated noise, Σ^*′*^(−*t*) = Σ^*′*^(*t*)^T^ implies *κ*_*nm*_(−*t*) = *κ*_*mn*_(*t*).

### Equal-time covariance Σ_ξ_

Inserting the decomposition (B2), the solution Eq. (31) of the continuous Lyapunov equation (29) reads

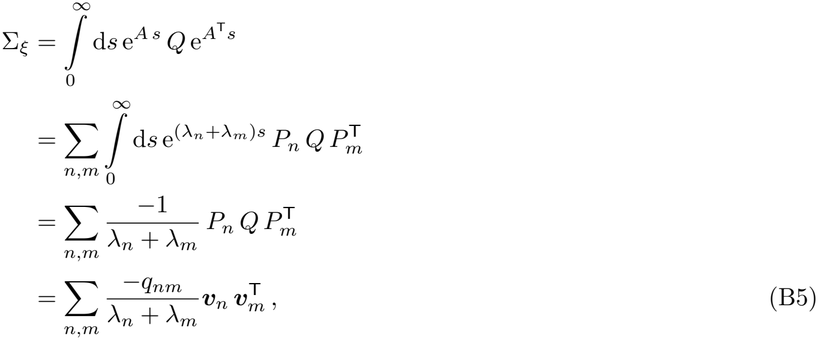

which yields the equal-time covariance Σ_*ξ*_ of ***x***(*t*) in the eigenvector basis of *W* with coefficients

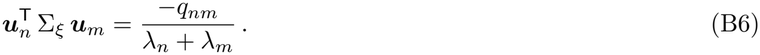

The non-diagonal matrix elements in Eq. (B5) describe the correlations between fluctuations along different eigenvectors ***v***_*n*_ and ***v***_*m*_ of *W* . They are proportional to *q*_*nm*_ in Eq. (B2), which depends only on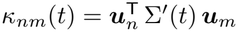. Thus, the effect of the recurrent network dynamics is to modulate the matrix elements in Eq. (B5) by dividing through the eigenvalues.

### Fisher information via spectral decomposition

Since for propagated (Gaussian) noise, the Fisher information (32) depends on Σ_*ξ*_ only via the term

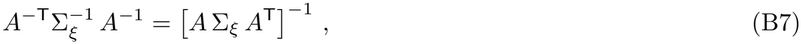

we compute

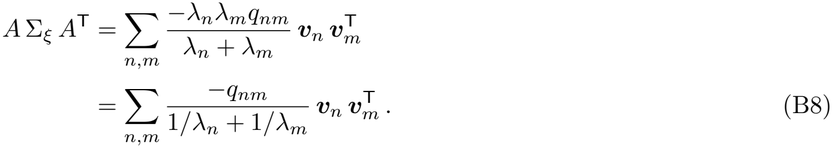

This symmetric, positive semi-definite matrix combines the effect of the recurrent connections on the neural manifold ***x***^*\**^(***z***) and on the (equal-time) noise covariance Σ_*ξ*_. It acts as an effective noise covariance matrix in the Fisher information written as if the recurrent connections are absent:

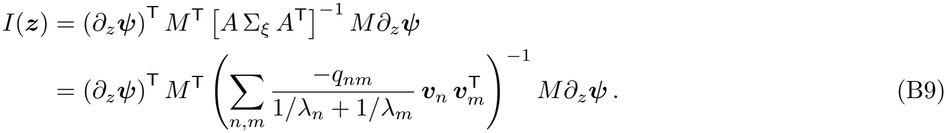

As discussed above, the effect of the recurrent network dynamics is to modulate the matrix elements in Eq. (B9) through the eigenvalues.

#### Slow and fast noise

For slowly varying noise, Σ^*′*^(*t*) ≈ Σ^*′*^(0) and thus *κ*_*nm*_(*t*) ≈ *κ*_*nm*_(0) = *κ*_*nm*_, and the integral in Eq. (B4) results in

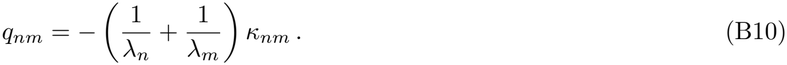

Inserting the coefficients *q*_*nm*_ given by Eq. (B10) into the effective covariance matrix Eq. (B9) in the Fisher information and using the decomposition Eq. (B2) yields

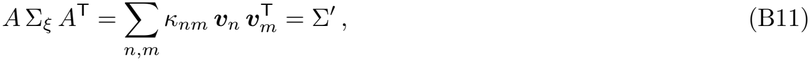

where Σ^*′*^ = Σ^*′*^(0) is the covariance matrix of the slow noise. Thus, the effects of the recurrent weights *W* on the Fisher information cancel, which then no longer depends on *W* .

This does not hold for fast noise: For temporally white noise, 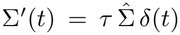 with noise strength covariance matrix 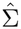 and the Dirac delta function *δ*(*t*), we have 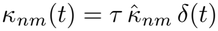 correspondingly. In this case, we simply get 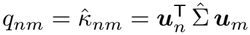. Consequently, in

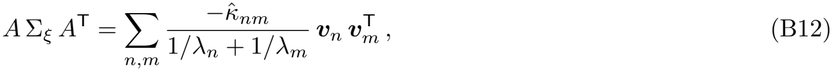

the modulation by the eigenvalues of *A* = *W* − 1 reflects the filtering of the noise combined with the amplification or attenuation of the recurrent stimulus representation.

## Appendix C: Scaled isometry of grid cell modules for lattice phase offsets

In this appendix, we derive the scaled isometry property of the population activity ***h***(***z***) = (*h*_1_(***z***), …, *h*_*N*_ (***z***))^T^ introduced in Sec. III C 1, with components 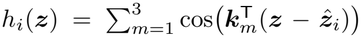 and spatial phase offsets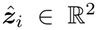.

Specifically, we show that for aligned hexagonal lattice configuration with *N* = *n*^2^ phases, the pullback metric is

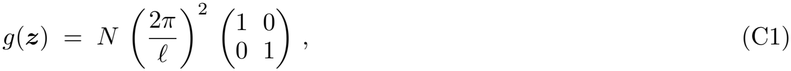

independently of ***z***, and we briefly sketch an extension to non-aligned (rotated) hexagonal lattice configurations.

### Fourier expansion of the pullback metric

Differentiating *h*_*i*_ gives 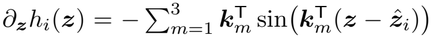, so the pullback metric (a 2 × 2 matrix) reads

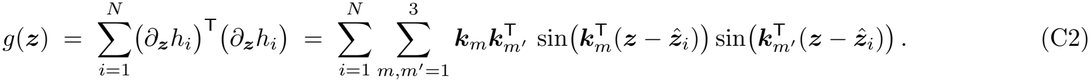

We expand the product of sines via sin *α* sin 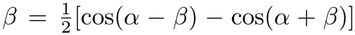 and write the ensuing sum over *i* as 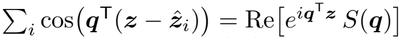 for the spatial frequencies ***q*** = ***k*** ± ***k*** with the *structure factor* of the phase offset configuration 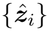 defined as

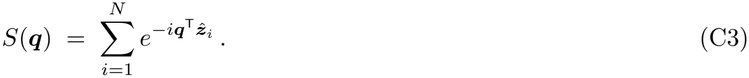

Then, the metric becomes a finite Fourier sum

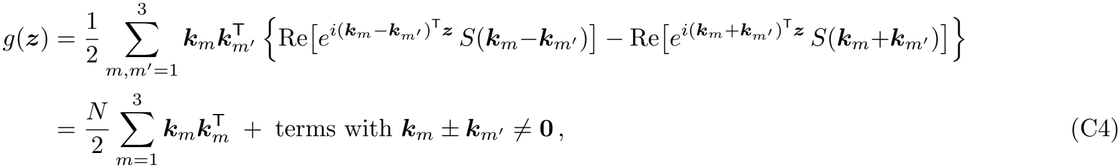

where the ***z***-independent part in the second line collects the *m*^*′*^ = *m* contribution to the difference terms (with *S*(**0**) = *N*). The nonzero frequencies ***k***_*m*_±***k***_*m*_^*′*^ give rise to ***z***-dependent cosine terms in the metric. Using ***k***_2_ = ***k***_1_ +***k***_3_, the occurring frequencies (wave vectors) are

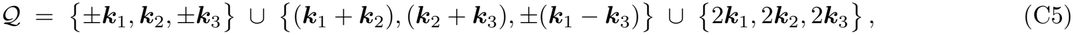

with squared lengths ∣***k***_*m*_ *∣*^2^, 3 ∣***k***_*m*_∣ ^2^ and 4 ∣***k***_*m*_ *∣*^2^, respectively. These 12 vectors lie in the first three shells of the reciprocal lattice *L*^*\**^ of the hexagonal lattice *L* formed by the grid pattern. The vectors ***k***_2_, ***k***_1_ + ***k***_2_ and ***k***_2_ + ***k***_3_ occur twice in the Fourier sum in Eq. (C4), accounting for the total of 15 terms with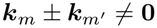.

The constant part in Eq. (C4) is already proportional to the identity matrix, as can be seen by explicit calculation (or by the sixfold symmetry of the wave vectors), 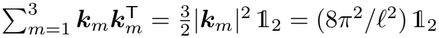, establishing the prefactor *N* (2*π/ℓ*)^2^ in Eq. (C1).

The scaled isometry property therefore reduces to the condition that the structure factor Eq. (C3) vanishes for every frequency vector in the set *Q*:

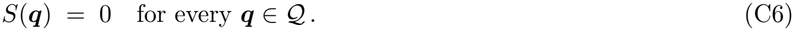

Since 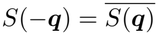 for real offsets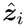, only nine representative vectors (one per ± -pair) need to be checked.

### Aligned hexagonal lattice with N = *n*^2^ phases

Let ***v***_1_, ***v***_2_ *∈* −^2^, |***v***_1_| = |***v***_2_|, generate the hexagonal lattice *L* of the grid pattern (so 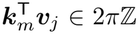 for all *m, j*). The *N* = *n*^2^ aligned lattice phase offsets that tile the parallelogram unit cell of *L* are given by

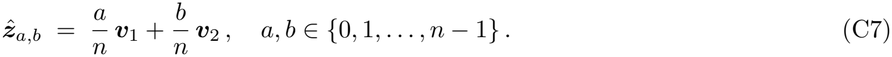

With these offsets, the structure factor reads

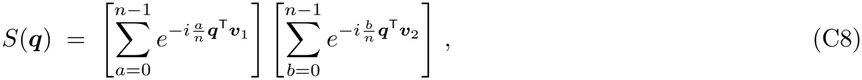

where each factor is a finite geometric sum:

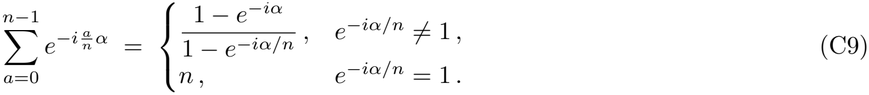

Since ***q*** ∈ *L*^*\**^ and ***v***_*j*_ ∈ *L*, we have *α* = ***q***^T^***v***_*j*_ ∈ 2*π*ℤ and *e*^*Δiα*^ = 1 for *j* = 1, 2. Consequently, the structure factor *S*(***q***) vanishes as long as *e*^*Δiα/n*^ ≠= 1 for at least one ***v***_*j*_. For ***q*** ∈ Q, the maximum absolute value of *α* = ***q***^T^***v***_*j*_ is 4*π*, obtained for ***q*** = 2***k***_*m*_. Hence, for *n*≥ 3, *e*^*Δiα/n*^ =≠ 1 (for at least one ***v***_*j*_) holds for every ***q*** ∈ Q, in other words Q ∩ *nL*^*\**^ = ∅. This proves the local isometry property and Eq. (C1) for *N* = *n*^2^∈ {9, 16, 25, … .}.

For *N* = 4 (*n* = 2), the frequency vectors 2***k***_*m*_ lie in 2*L*^*\**^, so the corresponding Fourier coefficients *S*(2***k***_*m*_) = *n*^2^ = 4 survive. The metric thus oscillates in ***z***; explicitly,

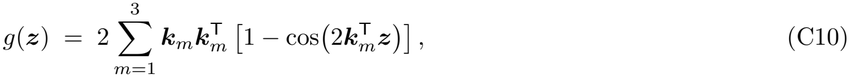

which degenerates (*g*(***z***) = **0**) at the points of 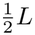, where all three 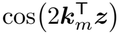 equal 1 simultaneously. Therefore, aligned configurations require *n* ≥ 3.

*Non-aligned hexagonal lattices*. The same conclusion (C1) holds whenever the offsets 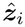 are taken from a more fine-grained hexagonal lattice *L*^*′*^⊃ *L* with *N* phases per unit cell of *L*, provided *N* ≥5.

This can be shown by applying a shift-invariance (“translation trick”) to the structure factor *S*(***q***): If we shift the offsets by any vector ***ϕ*** ∈ *L*^*′*^, then we obtain the equation

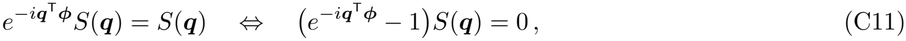

which holds for all ***q*** ∈*L*^*\**^, the reciprocal lattice of the hexagonal grid pattern lattice *L*, and in particular for every ***q*** ∈ *Q* . This equation can be understood as follows: Because the finer offset lattice *L*^*′*^ is periodic with respect to the (coarser) grid pattern lattice *L*, shifting the *N* offsets 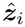 by any lattice vector ***ϕ*** ∈*L*^*′*^ just permutes them after folding them back to the unit cell of *L* using the modulo operation. Since the structure factor Eq. (C3) is a sum and a *L*-periodic function of the offsets 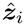 when ***q*** ∈ *L*^*\**^, *S*(***q***) remains unchanged, yielding Eq. (C11). Therefore, if ***q*** ∉*/* (*L*^*′*^)^*\**^, the reciprocal lattice of 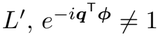 for at least one ***ϕ*** ∈ *L* ^*′*^, and the structure factor *S*(***q***) must be zero. The shortest nonzero vector of (*L*^*′*^)^*\**^ has length 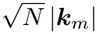, which exceeds 2|***k***_*m*_| as soon as *N* ≥ 5, hence *S*(***q***) vanishes on *Q* as required by condition (C6).

### Löschian hexagonal lattices

Hexagonal lattices *L*^*′*^ ⊃ *L* with *N* points per unit cell (i.e., per lattice point) of a coarser lattice *L* exist only for *N* ∈ℕ of the form *N* = *a*^2^ + *ab* + *b*^2^ with *a, b* ∈ ℤ (Löschian numbers [73, 74]). The reason for this is that the ratio of the distances between neighboring points in *L* and neighboring points in *L*^*′*^ is 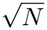 and that a vector ***v*** ∈ ℝ^2^ connecting neighboring lattice points of *L* must also be a lattice vector of the finer lattice *L*^*′*^. Specifically, with ***u***_1_, ***u***_2_ ∈ ℝ^2^, |***u***_1_| = |***u***_2_|, generating *L*^*′*^, we have ***v*** = *a* ***u***_1_ + *b* ***u***_2_ for *a, b* ∈ ℤ. With 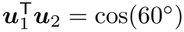, the squared distance between neighboring lattice points in *L* then yields

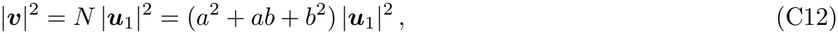

showing that *N* must be a Löschian number. The cases *N* = *a*^2^ are the aligned configurations considered above. For non-square *N*, the two lattices are not aligned and the rotation angle *α* of ***v*** relative to ***u***_1_ can be computed using tan 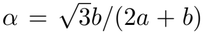. Together with *N* ≥ 5, the admissible numbers of hexagonal lattice phase offsets (numbers of grid cells in the module) are *N* ∈{7, 9, 12, 13, 16, 19, 21, 25, … }. In particular, the smallest admissible number is *N* = 7 with rotation angle arctan 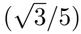 ≈ 19.1^*°*^. Schøyen *et al*. [35] numerically optimized the phase offsets to obtain a scaled (conformal) isometry and found that the minimal number is *N* = 7, realized by a hexagonal phase configuration that agrees with the one described here. In all cases, the pullback metric *g*(***z***) is given by Eq. (C1).

## Appendix D: Computation of the Ricci scalar

As a measure of intrinsic curvature, we consider the Ricci scalar curvature *R*, which derives from a contraction of the Riemann curvature tensor:

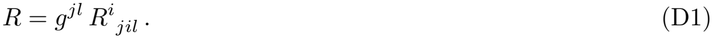

The Riemann curvature tensor

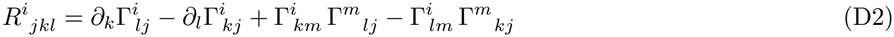

itself depends on the metric tensor *g*_*mk*_ (up to second order derivatives) via the Christoffel symbols

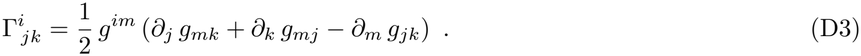

To simplify notation, we used here the Einstein sum convention (sum over the same upper and lower indices), *g*^*im*^ for the matrix elements of the inverse metric, and 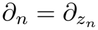.

For the grid cell network in Sec. III C 2, the Riemann curvature tensor can be explicitly computed since metric, Christoffel symbols and all their derivatives only depend on derivatives

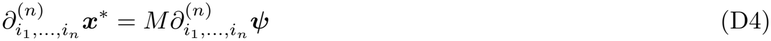

with

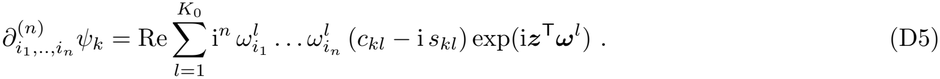

Moreover, derivatives of the inverse metric *g*^*im*^ can be explicitly expressed through 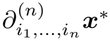 via

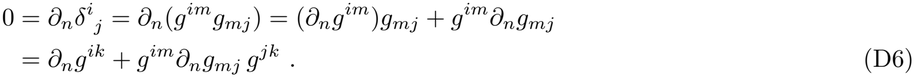

## References

[1] N. Brunel, J.-P. Nadal, and G. Toulouse, Information capacity of a perceptron, J. Phys. A: Math. Gen. 25, 5017 (1992).

[2] S. Chung, D. D. Lee, and H. Sompolinsky, Classification and geometry of general perceptual manifolds, Phys. Rev. X 8, 031003 (2018).

[3] H. Shan, Q. Li, and H. Sompolinsky, Order parameters and phase transitions of continual learning in deep neural networks, Proc. Natl. Acad. Sci. USA 123, e2501899123

[4] S. Bernardi, M. K. Benna, M. Rigotti, J. Munuera, S. Fusi, and C. D. Salzman, The geometry of abstraction in the hippocampus and prefrontal cortex, Cell 183, 954 (2020).

[5] J. Schmidhuber, Deep learning in neural networks: An overview, Neural Netw. 61, 85 (2015).

[6] Y. LeCun, Y. Bengio, and G. Hinton, Deep learning, Nature 521, 436 (2015).

[7] J. J. Jun, N. A. Steinmetz, J. H. Siegle, D. J. Denman, M. Bauza, B. Barbarits, A. K. Lee, C. A. Anastassiou, A. Andrei, Ç. Aydin, et al., Fully integrated silicon probes for high-density recording of neural activity, Nature 551, 232 (2017).

[8] S. Aimon, T. Katsuki, T. Jia, L. Grosenick, M. Broxton, K. Deisseroth, T. J. Sejnowski, and R. J. Greenspan, Fast near-whole-brain imaging in adult Drosophila during responses to stimuli and behavior, PLoS Biol. 17, e2006732 (2019).

[9] J. A. Gallego, M. G. Perich, L. E. Miller, and S. A. Solla, Neural manifolds for the control of movement, Neuron 94, 978 (2017).

[10] C. Langdon, M. Genkin, and T. A. Engel, A unifying perspective on neural manifolds and circuits for cognition, Nat. Rev. Neurosci. 24, 363 (2023).

[11] M. G. Perich, D. Narain, and J. A. Gallego, A neural manifold view of the brain, Nat. Neurosci. 28, 1582 (2025).

[12] C. Curto and N. Sanderson, Topological neuroscience: Linking circuits to function, Annu. Rev. Neurosci. 48, 491 (2025).

[13] R. Chaudhuri, B. Gerçek, B. Pandey, A. Peyrache, and I. Fiete, The intrinsic attractor manifold and population dynamics of a canonical cognitive circuit across waking and sleep, Nat. Neurosci. 22, 1512 (2019).

[14] R. J. Gardner, E. Hermansen, M. Pachitariu, Y. Burak, N. A. Baas, B. A. Dunn, M.-B. Moser, and E. I. Moser, Toroidal topology of population activity in grid cells, Nature 602, 123 (2022).

[15] N. Kriegeskorte and X.-X. Wei, Neural tuning and representational geometry, Nat. Rev. Neurosci. 22, 703 (2021).

[16] M. M. Monsalve-Mercado, G. M. Stine, M. N. Shadlen, and K. D. Miller, The geometry of the neural state space of decisions, bioRxiv 10.1101/2025.01.24.634806 (2025).

[17] C. E. Schoonover, S. N. Ohashi, R. Axel, and A. J. P. Fink, Representational drift in primary olfactory cortex, Nature 594, 541 (2021).

[18] S.-i. Amari, Information Geometry and Its Applications, Applied Mathematical Sciences, Vol. 194 (Springer, Tokyo, 2016).

[19] N. Ay, J. Jost, H. V. Lê, and L. Schwachhöfer, Information Geometry, Ergebnisse der Mathematik und ihrer Grenzgebiete, 3. Folge, Vol. 64 (Springer, Cham, 2017).

[20] J. Beck, V. R. Bejjanki, and A. Pouget, Insights from a simple expression for linear Fisher information in a recurrently connected population of spiking neurons, Neural Comput. 23, 1484 (2011).

[21] C. F. Gauss, General Investigations of Curved Surfaces of 1827 and 1825, translated by J. C. Morehead and A. M. Hiltebeitel (Princeton University Library, Princeton, NJ, 1902).

[22] W. M. Boothby, An Introduction to Differentiable Manifolds and Riemannian Geometry, Pure and Applied Mathematics (Academic Press, New York, 1975).

[23] D. J. Struik, Lectures on Classical Differential Geometry, 2nd ed. (Addison-Wesley, Reading, MA, 1961).

[24] A. Lingner, M. Pecka, C. Leibold, and B. Grothe, A novel concept for dynamic adjustment of auditory space, Sci. Rep. 8, 8335 (2018).

[25] E. L. Lehmann and G. Casella, Theory of Point Estimation, 2nd ed., Springer Texts in Statistics (Springer, New York, 1998).

[26] M. S. Bartlett, The square root transformation in analysis of variance, Supplement to the Journal of the Royal Statistical Society 3, 68 (1936).

[27] P. J. Antsaklis and A. N. Michel, Linear Systems (McGraw-Hill, New York, 1997).

[28] J. D. Semedo, A. Zandvakili, C. K. Machens, B. M. Yu, and A. Kohn, Cortical areas interact through a communication subspace, Neuron 102, 249 (2019).

[29] S. R. Jammalamadaka and A. SenGupta, Topics in Circular Statistics, Series on Multivariate Analysis, Vol. 5 (World Scientific, Singapore, 2001).

[30] H. S. Seung and H. Sompolinsky, Simple models for reading neuronal population codes, Proc. Natl. Acad. Sci. USA 90, 10749 (1993).

[31] M. Fyhn, S. Molden, M. P. Witter, E. I. Moser, and M.-B. Moser, Spatial representation in the entorhinal cortex, Science 305, 1258 (2004).

[32] T. Hafting, M. Fyhn, S. Molden, M.-B. Moser, and E. I. Moser, Microstructure of a spatial map in the entorhinal cortex, Nature 436, 801 (2005).

[33] Z. Ye and R. Wessel, Speed modulations in grid cell information geometry, Nat. Commun. 16, 7723 (2025).

[34] H. Stensola, T. Stensola, T. Solstad, K. Frøland, M.-B. Moser, and E. I. Moser, The entorhinal grid map is discretized, Nature 492, 72 (2012).

[35] V. S. Schøyen, K. Beshkov, M. B. Pettersen, E. Hermansen, K. Holzhausen, A. Malthe-Sørenssen, M. Fyhn, and M. E. Lepperød, Hexagons all the way down: Grid cells as a conformal isometric map of space, PLoS Comput. Biol. 21, e1012804 (2025).

[36] T. Solstad, E. I. Moser, and G. T. Einevoll, From grid cells to place cells: A mathematical model, Hippocampus 16, 1026 (2006).

[37] J. L. van Hemmen and C. Leibold, Elementary excitations of biomembranes: Differential geometry of undulations in elastic surfaces, Phys. Rep. 444, 51 (2007).

[38] E. Kropff and A. Treves, The emergence of grid cells: Intelligent design or just adaptation?, Hippocampus 18, 1256 (2008).

[39] A. Stepanyuk, Self-organization of grid fields under supervision of place cells in a neuron model with associative plasticity, Biol. Inspired Cogn. Archit. 13, 48 (2015).

[40] Y. Dordek, D. Soudry, R. Meir, and D. Derdikman, Extracting grid cell characteristics from place cell inputs using non-negative principal component analysis, eLife 5, e10094 (2016).

[41] M. M. Monsalve-Mercado and C. Leibold, Hippocampal spike-timing correlations lead to hexagonal grid fields, Phys. Rev. Lett. 119, 038101 (2017).

[42] T. D’Albis and R. Kempter, A single-cell spiking model for the origin of grid-cell patterns, PLoS Comput. Biol. 13, e1005782 (2017).

[43] S. N. Weber and H. Sprekeler, Learning place cells, grid cells and invariances with excitatory and inhibitory plasticity, eLife 7, e34560 (2018).

[44] A. Rahimi and B. Recht, Random features for large-scale kernel machines, in Advances in Neural Information Processing Systems, Vol. 20, edited by J. C. Platt, D. Koller, Y. Singer, and S. T. Roweis (Curran Associates, Red Hook, NY, 2007) pp. 1177–1184.

[45] L. D. Brown, T. T. Cai, and A. DasGupta, Interval estimation for a binomial proportion, Stat. Sci. 16, 101 (2001).

[46] J. B. Tenenbaum, V. de Silva, and J. C. Langford, A global geometric framework for nonlinear dimensionality reduction, Science 290, 2319 (2000).

[47] R. Ghrist, Barcodes: The persistent topology of data, Bull. Am. Math. Soc. 45, 61 (2008).

[48] U. Bauer, Ripser: Efficient computation of Vietoris–Rips persistence barcodes, J. Appl. Comput. sTopol. 5, 391 (2021).

[49] M. Poth and A. V. M. Herz, Burst activity plays no role in the field-to-field variability and rate remapping of grid cells, Hippocampus 31, 1128 (2021).

[50] A. Chadwick, A. G. Khan, J. Poort, A. Blot, S. B. Hofer, T. D. Mrsic-Flogel, and M. Sahani, Learning shapes cortical dynamics to enhance integration of relevant sensory input, Neuron 111, 106 (2023).

[51] M. C. Fuhs and D. S. Touretzky, A spin glass model of path integration in rat medial entorhinal cortex, J. Neurosci. 26, 4266 (2006).

[52] Y. Burak and I. R. Fiete, Accurate path integration in continuous attractor network models of grid cells, PLoS Comput. Biol. 5, e1000291 (2009).

[53] B. Lin and N. Kriegeskorte, The topology and geometry of neural representations, Proc. Natl. Acad. Sci. USA 121, e2317881121 (2024).

[54] M. M. Monsalve-Mercado and C. Leibold, Effect of boundaries on grid cell patterns, Phys. Rev. Res. 2, 043137 (2020).

[55] K. M. Gothard, W. E. Skaggs, K. M. Moore, and B. L. McNaughton, Binding of hippocampal CA1 neural activity to multiple reference frames in a landmark-based navigation task, J. Neurosci. 16, 823 (1996).

[56] J. L. Gauthier and D. W. Tank, A dedicated population for reward coding in the hippocampus, Neuron 99, 179 (2018).

[57] C. Tessereau, F. Xuan, J. R. Mellor, P. Dayan, and D. Dombeck, Navigating uncertainty: Reward location variability induces reorganization of hippocampal spatial representations, bioRxiv 10.1101/2025.01.06.631465 (2025).

[58] F. Acosta, S. Sanborn, K. Dao Duc, M. Madhav, and N. Miolane, Quantifying extrinsic curvature in neural manifolds, in Proceedings of the IEEE/CVF Conference on Computer Vision and Pattern Recognition Workshops (2023) pp. 610–619.

[59] M. Hauser and A. Ray, Principles of Riemannian geometry in neural networks, in Advances in Neural Information Processing Systems, Vol. 30, edited by I. Guyon, U. von Luxburg, S. Bengio, H. Wallach, R. Fergus, S. Vishwanathan, and R. Garnett (Curran Associates, Red Hook, NY, 2017) pp. 2807–2816.

[60] B. S. Robinson, R. Norman-Tenazas, M. Cervantes, D. Symonette, E. C. Johnson, J. Joyce, P. K. Rivlin, G. M. Hwang, K. Zhang, and W. Gray-Roncal, Online learning for orientation estimation during translation in an insect ring attractor network, Sci. Rep. 12, 3210 (2022).

[61] P. Vafidis, D. Owald, T. D’Albis, and R. Kempter, Learning accurate path integration in ring attractor models of the head direction system, eLife 11, e69841 (2022).

[62] F. Mastrogiuseppe and S. Ostojic, Linking connectivity, dynamics, and computations in low-rank recurrent neural networks, Neuron 99, 609 (2018).

[63] L. Pezon, V. Schmutz, and W. Gerstner, Linking neural manifolds to circuit structure in recurrent networks, Neuron 114, 1682 (2026).

[64] J. J. Hopfield, Neural networks and physical systems with emergent collective computational abilities, Proc. Natl. Acad. Sci. USA 79, 2554 (1982).

[65] H. Sompolinsky and I. Kanter, Temporal association in asymmetric neural networks, Phys. Rev. Lett. 57, 2861 (1986).

[66] J. Buhmann and K. Schulten, Storing sequences of biased patterns in neural networks with stochastic dynamics, in Neural Computers, edited by R. Eckmiller and C. von der Malsburg (Springer, Berlin, Heidelberg, 1989) pp. 231–242.

[67] A. V. M. Herz, Z. Li, and J. L. van Hemmen, Statistical mechanics of temporal association in neural networks with transmission delays, Phys. Rev. Lett. 66, 1370 (1991).

[68] C. Leibold and R. Kempter, Memory capacity for sequences in a recurrent network with biological constraints, Neural Comput. 18, 904 (2006).

[69] C. Leibold, A model for navigation in unknown environments based on a reservoir of hippocampal sequences, Neural Netw. 124, 328 (2020).

[70] X.-X. Lin, Y.-H. Yiu, and C. Leibold, Emergence of spatial representation in an actor-critic agent with hippocampus-inspired sequence generator, in International Conference on Learning Representations (2026) arXiv:2510.09951 [q-bio.NC].

[71] A. Pellegrino and A. Chadwick, RNNs perform task computations by dynamically warping neural representations, in Advances in Neural Information Processing Systems, Vol. 38 (Neural Information Processing Systems Foundation, San Diego, CA, 2025) pp. 4003–4036.

[72] A. A. Russo, S. R. Bittner, S. M. Perkins, J. S. Seely, B. M. London, A. H. Lara, A. Miri, N. J. Marshall, A. Kohn, T. M. Jessell, et al., Motor cortex embeds muscle-like commands in an untangled population response, Neuron 97, 953 (2018).

[73] A. Lösch, The Economics of Location, translated by W. H. Woglom with the assistance of W. F. Stolper (Yale University Press, New Haven, CT, 1954).

[74] OEIS Foundation Inc., Sequence A003136 (Löschian numbers), The On-Line Encyclopedia of Integer Sequences, https://oeis.org/A003136 (2026), accessed s7 May 2026.

